# Development and assessment of fully automated and globally transitive geometric morphometric methods, with application to a biological comparative dataset with high interspecific variation

**DOI:** 10.1101/086280

**Authors:** Tingran Gao, Gabriel S. Yapuncich, Ingrid Daubechies, Sayan Mukherjee, Doug M. Boyer

## Abstract

Automated geometric morphometric methods are promising tools for shape analysis in comparative biology: they improve researchers’ abilities to quantify biological variation extensively (by permitting more specimens to be analyzed) and intensively (by characterizing shapes with greater fidelity). Although use of these methods has increased, automated methods have some notable limitations: pairwise correspondences are frequently inaccurate or lack transitivity (i.e., they are not defined with reference to the full sample). In this study, we reassess the accuracy of two previously published automated methods, cPDist [1] and auto3Dgm [2], and evaluate several modifications to these methods. We show that a substantial fraction of alignments and pairwise maps between specimens of highly dissimilar geometries were inaccurate in the study of Boyer et al. [1], despite a taxonomically sensitive variance structure of continuous Procrustes distances. We also show these inaccuracies can be remedied by utilizing a *globally informed* methodology within a collection of shapes, instead of only comparing shapes in a pairwise manner (c.f. [2]). Unfortunately, while global information generally enhances maps between dissimilar objects, it can degrade the quality of correspondences between similar objects due to the accumulation of numerical error. We explore a number of approaches to mitigate this degradation, quantify the performance of these approaches, and compare the generated pairwise maps (as well as the shape space characterized by these maps) to a “ground truth” obtained from landmarks manually collected by geometric morphometricians. Novel methods both improve the quality of the pairwise correspondences relative to cPDist, and achieve a taxonomic distinctiveness comparable to auto3Dgm.

## Introduction

Quantifying and comparing complex shapes is a key component of fields as diverse as evolutionary morphology, molecular biochemistry, computer vision, and computational anatomy. A variety of analytical methods have been developed to achieve this goal, including landmark-based geometric morphometrics [3–6], voxel-based morphometry [7], and spherical harmonics [8–9]. Of these methods, three-dimensional geometric morphometrics (3DGM) based on the alignment of spatial coordinates through Procrustes superimposition is particularly widespread in evolutionary morphological studies (for reviews, see [10–12]). Though popular, 3DGM is nonetheless a time-consuming and labor-intensive process, requiring a substantial number of landmarks to be placed on each specimen by the researcher [13–15]. The simulation study of [16] suggested shape characterization is unstable without at least 30 landmarks, a number which may not be feasible in samples spanning multiple genera (e.g., [1]). For researchers utilizing 3DGM, the reliance on user-determined landmarks generates a trade-off between sample size and detail of morphological representation for a given time spent collecting data, and thus limits the explanatory power of morphological data. Without significant methodological advances, future geometric morphometric morphological studies are likely to remain limited.

In order to more thoroughly characterize shape variation and decrease processing time, geometric morphometric approaches have become increasingly automated, including both semiautomated (based on semilandmarks [17–22] or eigensurfaces [13,23]) and fully automated [1,2,24,25] shape characterization methods. Automated 3DGM methods improve researchers’ ability to sample phenotypes intensively (by increasing the resolution of shape characterization) and extensively (by permitting the inclusion of more specimens). These outcomes neatly align with Houle et al.’s [26] recommendations for advancing phenomics – the study of high-dimensional phenotypic data.

Initial evaluations of automated 3DGM methods recover performance similar to or better than user-determined landmarks for species discrimination [1,2] and shape characterization of certain types of shapes [27]. Still, as with user-based approaches, current automated 3DGM methods suffer from limitations. The first limitation concerns sample availability: a significant investment of time is required to convert specimens into 3D digital models. This limitation will be reduced as researchers continue to contribute data to online repositories such as MorphoSource [28]. The second limitation concerns the analytical workflow: while the fully automated method published in [1], the continuous Procustes distance method (cPDist), successfully classifies specimens by species, the output of the method cannot be analyzed in a way analogous to user-determined landmarks due to the lack of *transitivity* of the resulting pairwise maps (i.e., the direct map from A to C is not the same as the map from A to B to C). Because automated landmarks are defined only on a pairwise basis, rather than the whole collection of samples, cPDist does not produce a set of globally consistent landmarks as required for downstream 3DGM analyses. The third limitation stems from the computational intensity of automated 3DGM methods: the method published in [2], auto3Dgm, produces a transitive set of “pseudolandmarks” that is applicable to the entire collection of samples, but the method does not begin to yield consistent results unless over 1,000 pseudolandmarks are identified on each specimen (at least when each specimen is discretized as a mesh of ~5,000 vertices) [29]. Even with a relatively powerful computer, auto3Dgm may take weeks to analyze a dataset of 200 specimens without access to parallel computational resources.

Despite these limitations, initial results and applications of automated 3DGM methods are encouraging [1,2,24,27,30,31]. Still, the ability of fully automated methods to achieve certain goals in biological research has not been thoroughly explored. This study addresses the key issues concerning previously published automated 3DGM methods [1,2], including:

1. Assessing error rates of cPDist and identifying general properties of shapes which may indicate whether automated mappings are likely to be accurate;
2. Evaluating the ability of an MST-based approach (cPMST, a variant of cPDist inspired by auto3Dgm) to avoid bad alignments;
3. Describing quantitative and qualitative differences in ordinated shape spaces recovered by automated 3DGM methods.

In addition to examining previously published automated methods, we also develop and evaluate several approaches for transitively joining dissimilar shapes with intermediate shape sequences. Since these approaches utilize global geometric information in the entire shape collection, we refer to these novel methods as “globally informed methods”. To increase ease of application of the methods introduced in this study, MATLAB code is provided for each method in the supporting information.

## Background on previous and related automated 3DGM methods

The cPDist method [1,24] begins by flattening disc-type shapes (such as a tooth crown without roots) into planar discs using a conformal (angle-preserving) projection. The algorithm then exhaustively searches the space of all conformal maps between the flattened shapes for an “optimal conformal map” minimizing an energy functional. Conformal maps between discs are characterized by the choice of a pair of correspondence points and an in-plane rotation (thus only three degrees of freedom), which makes the exhaustive search highly efficient. The resulting conformal map then serves as an initialization for a final thin plate spline (TPS) procedure, which strives to stretch the two flattened shapes so that regions of “high curvature” (e.g., cusp tips or tuberosities) align. The value of the energy functional on the final map (composition of TPS and the optimal conformal map) defines a distance between the pair of shapes, which we refer to as the continuous Procrustes (cP) distance [32]. When the cP distance is small, the final map is usually of high quality and can be leveraged to reveal interspecific variation (cf. [1]).

As an initial assessment of the biological relevance of these correspondence maps, Boyer et al. [1] compared the classification success rates of cPDist and Procrustes distances computed from user-determined landmarks (a more traditional morphometric approach) on a mammalian molar dataset. The comparison showed that cPDist was able to taxonomically classify specimens at a rate better than or equal to the method based on user-determined landmarks. Since then, Boyer et al. [24] used cPDist to confirm the attribution of a newly discovered fossil to a species previously believed to be much younger than indicated by dating of the fossil. However, due to lack of transitivity, it is not clear how the resulting correspondence maps of cPDist pipeline should be incorporated in a more traditional geometric morphometric workflow. Furthermore, when inspecting pairwise correspondence maps between specimens in detail, Boyer et al. [24] observed anomalies in some of the maps (e.g., reversed alignments of the buccolingual axis). Though the distance matrices and ordinations based on them produced intelligible results, these errors raised questions about possible inaccuracies lurking in the analysis. We examine these inaccuracies in more detail here.

Boyer et al. [2] reported a different automated method, auto3Dgm, which guarantees transitivity, thereby permitting more familiar modes of downstream analysis. First, auto3Dgm computes all pairwise alignments and distances with a modified version of the Iterative Closest Points (ICP) algorithm [33]. Transitivity is then imposed with the following procedure based on a Minimum Spanning Tree (MST) for the entire collection: 1) view the collection of shapes as a complete weighted graph (in which edge weights are defined by the pairwise distances), and extract an MST for this graph; 2) for any pair of shapes, the alignment between them is obtained by identifying the unique shortest path connecting them in the MST and composing the pairwise alignments along the edges constituting this path. By the nature of the MST, only alignments that yield small pairwise distances are involved in the final alignments. Auto3Dgm outputs a “pseudolandmark file” that can be analyzed with standard geometric morphometric software such as *morphologika*^2^ [34] or MorphoJ [35]. Boyer et al. [2] argued that this procedure should generally reduce the mapping errors, and verified this claim with three osteological datasets. R code for auto3Dgm is available based on its original MATLAB implementation [36].

Recently, Koehl and Haas [25] suggested minimizing a novel metric, the symmetric deformation energy, when searching for a globally optimal conformal map between two closed surfaces. Based on their analysis, the program MatchSurface outperforms cPDist in approximating the ground truth pairwise distances computed from user-determined landmarks, in correctly classifying specimens to taxonomic groups, and in generating phenetic trees that more closely resemble trees generated from user-determined landmarks. However, MatchSurface may suffer from similar problems we have noted with cPDist (i.e., potential anomalies in pairwise maps, lack of transitivity, inability to utilize the method in a traditional morphometric workflow). In addition, because user-based approaches such as 3DGM have their own methodological challenges, it is unclear if automated 3DGM methods should be evaluated primarily by their ability to replicate pairwise distances computed from user-determined landmarks (although this is also the approach we use in this study). An in-depth comparative analysis of MatchSurface and the methods introduced here is beyond the scope of this paper, but will be important for further development of automated geometric morphometric methods.

In a broader context, the development of automated 3DGM methods resonates with the emerging interest in the analysis of collections of shapes in the computer graphics community (e.g., [37–40], among others). The starting point is the observation that pairwise shape registration often yields more accurate results between pairs of similar shapes than dissimilar ones: the resulting pairwise correspondence maps are much more meaningful when the shapes are near-isometric. Therefore, when working with a large collection of shapes, one can usually find a sequence of (pairwise similar) intermediate shapes between any pair of dissimilar shapes, and build the correspondence map between them by composing the more accurate pairwise correspondences along this sequence. This composition strategy leads to more accurate maps than direct pairwise comparisons. The novel methods presented in this paper are all derived from this general idea, leveraging the size of the dataset and high-quality maps between similar shapes to improve the accuracy of correspondence maps between dissimilar shapes.

## Methods and materials

We assess the qualities of automated correspondence analyses using the dataset originally published in Boyer et al. [1], and available through various sources [41–43]. The sample consists of 116 mandibular second molars of living and fossil primates and their close relatives. Further details (such as included species and specimen information) can be found in the supplementary data of Boyer et al. [1]. Our main approach is to compare automated results of anatomical correspondence and geometric similarity among shapes in this dataset to a “ground truth” – a set of 16 user-determined landmarks placed on each specimen by experienced geometric morphometricians. As noted above, we have reservations about regarding user-determined landmarks as the performance standard for automated correspondence analyses, but this framework facilitates comparisons between user-based and automated approaches.

## Quantifying errors of cPDist

The accuracy of pairwise correspondences can be quantified using Mean Square Error (MSE). MSE is calculated by first mapping user-determined landmarks from one tooth to another using the automated pairwise correspondence (creating a set of “propagated landmarks” on the second tooth), and then taking the average of the squared Euclidean distances between the user-determined and propagated landmarks. Larger MSEs indicate greater deviations between the two sets of landmarks (and a greater discordance between the user-determined and automated assessments). While there are other approaches to assess accuracy relative to user-determined landmarks (including directly comparing distance matrices generated by each method or the ordinations resulting from those matrices), MSE benefits by focusing on the “local” inaccuracy of correspondence maps at the level of individual landmarks.

Boyer et al. [1] assessed cPDist’s error rate by computing a variant of MSE between user-determined and propagated landmarks (Supplementary Table 8 of [1]), which realigned propagated and user-determined landmarks with an additional Procrustes superimposition. Unfortunately, this realignment could potentially mask several types of mapping errors (Fig 1). Most notably, the cPDist algorithm may result in an axial inversion in which incorrect sides are matched to one another (buccal-lingual [Fig 1c] or anterior-posterior [Fig 1d] inversions). To assess the prevalence of mapping errors in the analysis of Boyer et al. [1], we perform an extensive (though not exhaustive) visual check of 16 propagated landmarks in 588 pairwise mappings. We observed five types of errors; the four primary errors are shown in Fig 1 (the fifth error type, a 90 ° rotation, occurred in only three of 161 instances of error). To test whether errors occur between dissimilar teeth, we compare the pairwise cP distances observed in the good and bad maps of this test sample.

**Fig 1.**
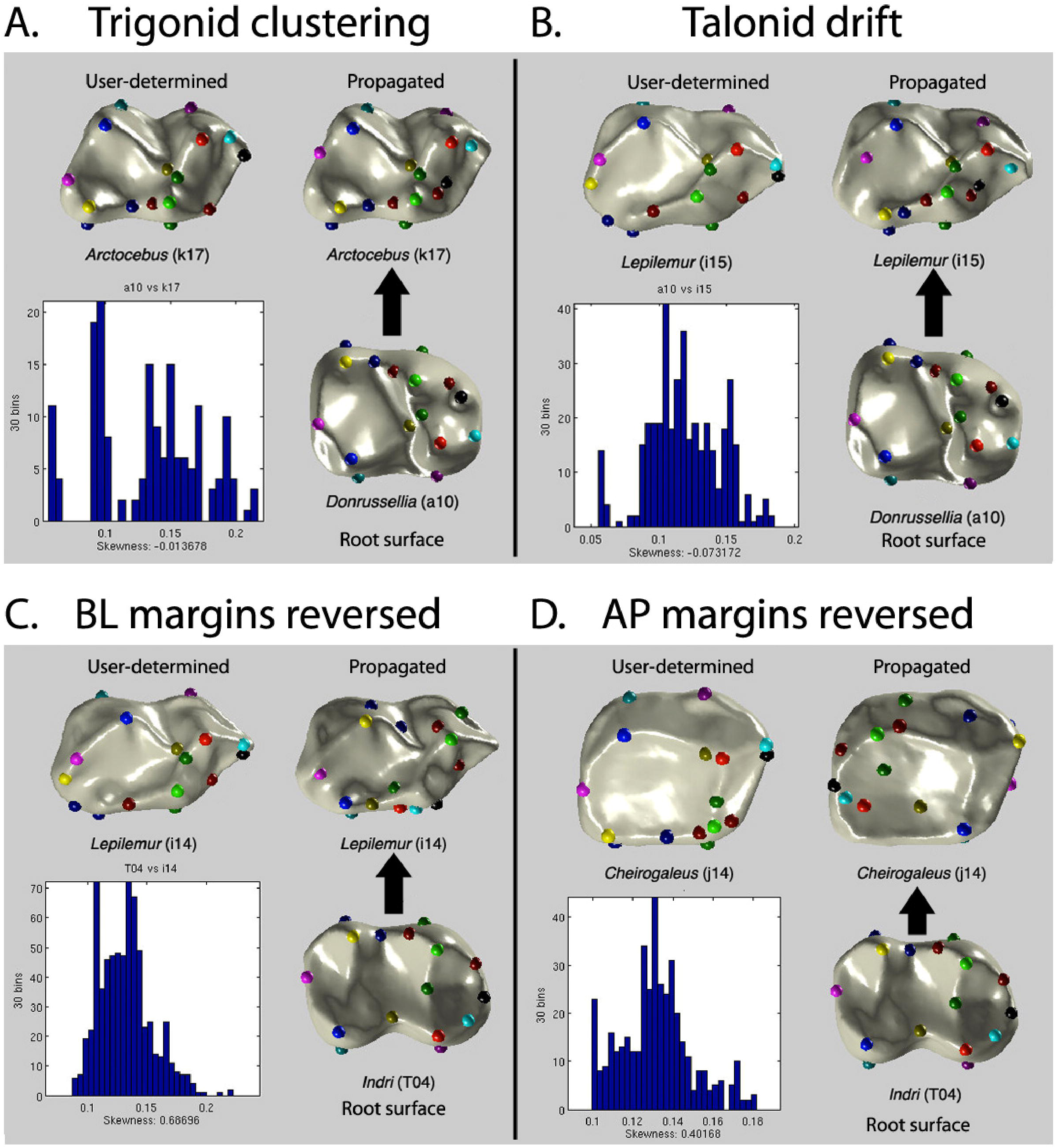
Classification of errors made by cPDist using MATLAB interface. Each panel shows the root surface with user-determined landmarks (lower right), user-determined landmarks on target surface (upper left), and propagated landmarks on target surface (upper right). The distribution of the values of the continuous Procrustes functional of candidate maps are shown in the lower left of each panel. The four main error types are pictured here and include A) Trigonid clustering: landmarks bunched in anterior portion of tooth; B) Talonid drift: landmarks spread over posterior portion of tooth; C) buccal-lingual (BL) inversion: landmarks are reversed from side-to-side; and D) anterior-posterior (AP) inversion: landmarks are reversed from front-to-back. A fifth error type, 90° rotation, is not shown, but was only observed three times.

We also test the hypothesis that more dissimilar teeth generate erroneous maps by comparing their Dirichlet Normal Energy (DNE), a measure of the bending energy of a surface that has been shown to partition species by diet reliably [44–46]. For a reduced sample of comparisons between specimens from extant species (n=138), species mean DNE values are taken from [44], and the absolute value of the difference between the source and target teeth calculated. Poor maps are expected to occur between teeth that exhibit a large difference in DNE.

Finally, we examine the skewness of the distribution of candidate maps for good and poor maps. In order to find an optimal map with minimum energy, cPDist searches among a large number (typically tens of thousands) of candidate conformal maps between two surfaces. In general, we expect good maps to be more strongly distinguished from the population of candidate maps than poor maps, as there are often many candidate maps with comparably low energy when a poor map forms the minimum in the exhaustive search. By measuring the skewness of the energy distribution of candidate maps, we assess the distinctiveness of the optimal map: if the optimal map is less distinct, the histogram of candidate maps should be skewed to the right (positive skew), forcing cPDist to select from several candidate maps with similar cP distances.

## Evaluating accuracy of a minimum-spanning tree approach

As described above, auto3Dgm [2] improves alignments between dissimilar shapes and imposes transitivity for the entire collection using MST. However, while MST improves pairwise cP maps, the quality of the resulting maps is not guaranteed. The visually inspected test sample permits the identification of a threshold cP distance below which all maps are good maps. To gauge the accuracy of the MST approach, we evaluate if any edge lengths in the MST exceed this threshold.

We also compare the landmark MSEs of cPDist to cPMST, another MST-based approach inspired by auto3Dgm [2]. While auto3Dgm’s representation of surfaces through pseudolandmarks is analogous to user-determined landmarks (increasing the method’s utility to comparative morphologists), these pseudolandmarks are not globally consistent in the same manner as user-determined landmarks. In auto3Dgm, pseudolandmarks are randomly sampled on each surface in a collection, without knowledge of the exact sampling procedure for other surfaces. The full set of user-determined landmarks is likely to be excluded from the set of pseudolandmarks, making it impossible to calculate landmark MSEs for auto3Dgm. Here, we evaluate the accuracy of cPMST, an upgraded version of cPDist that generates globally consistent maps between all pairs of shapes within a collection, motivated by the MST approach first adopted in auto3Dgm. With cPMST, pairwise maps are defined for the pair of surfaces in their entirety (and necessarily including user-determined landmarks), so that landmark MSEs can be computed.

## Development and refinement of globally informed methods

Though using an MST improves pairwise correspondences between dissimilar shapes, this approach could potentially degrade the quality of maps between similar shapes. Directly aligning similar teeth avoids the accumulation of random errors in pairwise comparisons, while composing alignments along a path through the MST involves intermediate shapes and can amplify random error. MSTs minimize the *total* sum of edge lengths in a subgraph connecting all the vertices; this global minimization permits large distortion of local distances. Many intermediate vertices may separate a pair of reasonably close vertices, so that the sum of the length of all intermediate hops may be larger than the direct pairwise distance.

The potential for map degradation with MST-based approaches and the results of our accuracy analyses (see Results) made it clear that there was ample room for improving previously published automated 3DGM methods. We propose several approaches that maintain global transitivity but also attempt to reduce the potential for local distance distortion. These approaches are based on the research of tree-based metric space approximation and dynamic programming in computer science. We examine the effects of several different methodological variations: 1) alternative methods to avoid map degradation, 2) alternative root shapes for landmark propagation when transforming maps into pseudolandmarks, 3) alternative methods for post-processing maps, and 4) the effects of pseudolandmark sampling resolution. All examined methods are summarized in Table 1 and described in detail below.

**Table 1.**
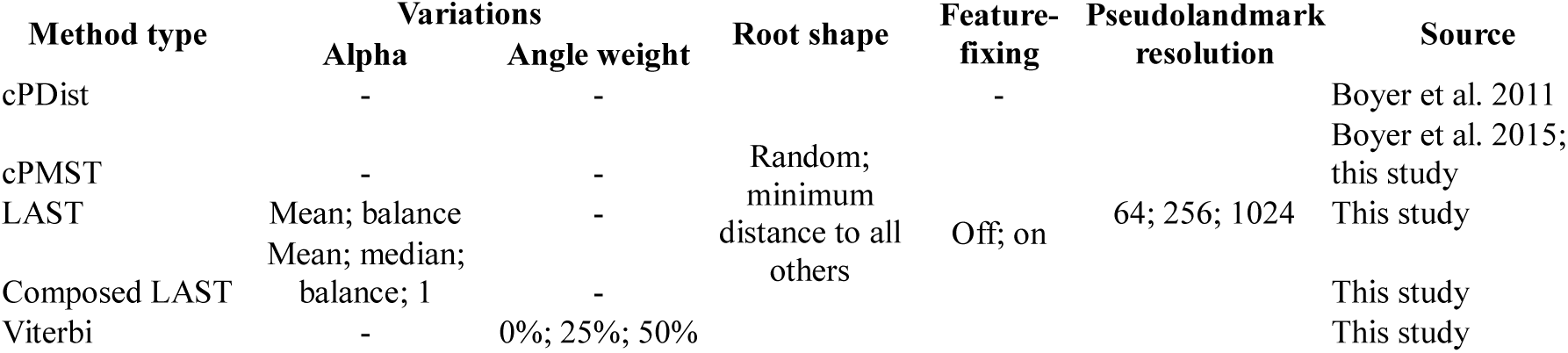
Globally informed methods developed for and analyzed in the current study.

### Alternative methods to avoid map degradation

Our first attempt to avoid potential degradation when mapping through intermediate shapes is to restrict the depth of the minimum spanning tree, and impose that any pair of shapes in the MST be separated by a controlled number (no greater than twice the tree depth) of intermediate shapes. This type of graph theoretic construction, known as a bounded-hop MST problem, is nondeterministic polynomial-time hard and has no practical polynomial-time and constant-factor approximation in general [47]. However, rather than globally minimizing the sum of edge lengths, it is almost trivial to minimize the number of hops along the tree that separates any pair of vertices: the best strategy is to designate one vertex as the root and connect any other vertices to it. Generalizing from counting the number of hops to measuring the length of the path, researchers study shortest-path trees (SPT) of a graph *G*, which are trees that span the graph with the property that any path connecting a vertex to the root is also the shortest path in the graph *G* between that particular vertex and the root. SPTs strive to minimize local distance distortions without controlling the total edge length, while MSTs minimize the latter but sacrifice the former. Leveraging the advantages of both MSTs and SPTs, the concept of Light Approximate Shortest-Path Tree (LAST) was developed to balance local distance distortion and total edge length [48]. We use LAST to alleviate the quality degradation of correspondences between similar teeth in the cP distance framework. Because LAST is also a tree, global transitivity is achieved.

Unlike MST and SPT, which are calculated directly from the data, LAST depends on a parameter, α (*≥* 1), that controls the trade-off between the advantages of MST and SPT. When α is close to 1, the LAST becomes more like a SPT; as α approaches infinity, the LAST becomes more similar to an MST. We focus on experimenting with two candidates for α: the mean of the local distance distortions on an MST and the special value 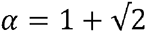, which generates a LAST “balanced” between the SPT and MST.

By altering how local distance distortion is computed, we develop a second set of LAST methods. Normally, computing the distance distortion of a tree between two vertices involves finding the shortest path on the tree that connects the pair, and then dividing the path length by the direct distance between the vertices. However, since the cP distance is defined as the minimum of an energy functional, a potentially more meaningful definition of local distance distortion uses the value of the energy functional obtained by composing pairwise correspondences along the shortest path (rather than the cumulative path length) in the numerator. We evaluate “composed” LAST methods with four candidates for α: the mean and median of the composed local distance distortions, the special value 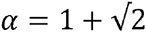, and α = 1 (which does not generate an SPT with composed local distance distortion^1^).

We also attempted to fix map degradation with an approach that does not rely on trees (though this method is not strictly transitive). For any pair of shapes, the goal is to find the best path between them such that the composition of pairwise correspondences along the hops gives the most meaningful map. In the spirit of cP distance, which produces a map that minimizes an energy functional, we use the value of the same energy functional (called the “cP value”) as an indicator for the quality of pairwise correspondences. Though the number of shapes in a collection is finite, the set of possible paths connecting a pair of shapes is exponentially large, making it difficult to search exhaustively for the optimum; computational complexity is even larger when looking for optimal paths between *all* pairs of shapes. This computational difficulty is tackled with a dynamical programming algorithm motivated by the Viterbi algorithm in the context of hidden Markov models. Each shape, viewed as a vertex in the complete distance graph, is treated as a hidden state of a Markov chain, and the transition probability from one state to another is determined by the cP distance between the two shapes, with smaller distances indicating greater probability. In this setup, the optimal path of composition between a pair of shapes can be interpreted as the most likely path connecting the two states. When the length of the path is fixed, optimal path searching can be done using the Viterbi algorithm (assuming the hidden state is identical to the observed state).

Due to the efficiency of the Viterbi algorithm, it is easy to compute the optimal paths of all possible numbers of hops (ranging from 1 to one less than the total number of shapes in the collection), and to choose the path along which the composition of pairwise correspondences leads to the lowest possible cP value. This method is denoted as “Viterbi”. Since direct links between two shapes are the only paths with 1 hop, the Viterbi method is guaranteed to keep direct pairwise maps if they produce the lowest cP values among all paths of different number of hops, thus avoiding the accumulation of random errors through propagation along long paths in an MST. Though global transitivity is not maintained, the determination of the unique optimal path connecting any two shapes leads to good correspondence maps.

For the Viterbi method, the transition probability between any two states can depend on metric geometric information other than path distance, such as the angle between consecutive hops in a path. More obtuse angles (i.e., closer to 180 degrees) between consecutive hops are likely preferable, as more acute angles lead to more torturous paths, which may increase random error accumulation. After proper renormalization, distances and angles can be combined with convex weights to determine the transition probability. In this study, we compare the effect of angles weighted at 0%, 25%, and 50% in the computation of transition probability.

### Alternative root shapes for pseudolandmark propagation

For these novel methods, globally consistent pseudolandmarks can be generated by randomly sampling a number of vertices and then propagating these pseudolandmarks to all remaining shapes after pairwise correspondences are established. These pseudolandmarks depend on the distribution of points sampled on the initial shape, but are independent of the choice of the initial shape (although since Viterbi methods are not strictly globally transitive, the initial shape does affect pseudolandmarks). To evaluate the effect of the choice of root shape on shape characterization, we generate pseudolandmarks from two different root shapes (a randomly chosen tooth [*Chronolestes simul* IVPP V10696-2] and the tooth with minimum average distance from other teeth in the collection) for all methods.

### Alternative methods of post-processing

The final novel development examined here is a post-processing step that performs an additional TPS procedure to align geometric characteristics (e.g., vertices of locally maximum conformal factors, locally maximum/minimum Gaussian curvatures) of two shapes. Since these geometric characteristics may be generally understood as “features”, we call this step “Feature-Fix”. Feature-fixing is similar to the last step in the computation of cP distances in Boyer et al. [1], which helps to correct the random errors accumulated through correspondence compositions but sometimes introduces artificiality in regions without geometric characteristics. We implement all novel methods with and without feature-fixing.

### Sampling density of pseudolandmarks

Several previous studies have highlighted the importance of landmark sampling density for shape characterization [16,29]. Vitek et al. [29] suggested that auto3Dgm does not produce consistent results unless 1000 pseudolandmarks are generated on each specimen. To examine the influence of sampling density on methods developed in this study, we implement all methods with 64, 256, and 1024 pseudolandmarks sampled.

### Comparing effects of globally informed methods on the characterization of geometric affinities

The different types of globally informed methods (n=8: MST, LAST, Viterbi and their variations), the root shape used to generate pseudolandmarks (n=2), the use of feature-fixing in post-processing (n=2), and the sampling density of pseudolandmarks (n=3) can be understood as “parameters” of the cP distance improvement framework. To compare variance patterns within and among these parameters, we first generate pseudolandmarks for the sample using all 120 distinct parameter combinations. Each combination can be represented by a scatterplot of 116 points (one for each tooth in the dataset) and compared using Procrustes analysis. To accomplish this, we reshape the x-y-z spatial coordinates of all pseudolandmarks on each tooth into a vector, and run principal component analysis (PCA) on the 116 vectors. The PCA generates 116 principal component scores for each of the 116 teeth in the dataset. Using the first three principal component scores as coordinates of the teeth creates a scatterplot in three-dimensional Euclidean space. This procedure essentially embeds an abstract metric structure into a Euclidean space of reduced dimensionality, to which a traditional landmark-based Procrustes analysis can be applied. Finally, we run generalized Procrustes analysis on these scatterplots in *morphologika*^*2.5*^ [34], and take the first 26 principal scores (the minimum number accounting for at least 95% of the variance) as a 26-dimensional feature vector encoding a parameter combination (S1 Appendix).

We implemented vector equivalents of one-way ANOVA, two-way ANOVA, and a linear mixed model in MATLAB (for further detail, see S2 Appendix). One-way ANOVA is used to evaluate the significance of each parameter. Two-way ANOVA reveals pairwise interaction effects among all parameters; we focus on interactions between “method” and each of the other parameters. The linear mixed model is used to determine which factors lead to shape characterizations that are most similar to the user-determined ground truth of the dataset.

### Evaluating accuracy of globally informed methods

We gauge the accuracy of the novel globally informed methods in two ways. First, we compare the landmark MSEs of all new methods to the landmark MSE generated by cPDist. The performance of cPMST (without feature-fixing) relative to cPDist was chose as a baseline for evaluating the accuracy of the globally informed methods presented here. A novel method is deemed preferable to the cPMST baseline if: 1) there are fewer pairwise mappings in which the landmark MSE of the novel method is greater than landmark MSE of cPDist (i.e., there are fewer positive residuals when the landmark MSEs of a novel method are plotted against the landmark MSEs of cPDist), or 2) the mean or maximum positive residual of a novel method is less than the mean or maximum positive residual of cPMST. Either of these patterns may indicate that the novel method does not accumulate random error through intermediate correspondences.

Comparing the distribution of landmark MSEs under each method provides another way to evaluate the accuracy of different methods. For inaccurate methods, the distribution of landmark MSEs may have a higher mean or display greater variance (or both) than the landmark MSE distribution of a more accurate method. Here we compare the landmark MSE distance matrices of all main methods using a Multiple Response Permutation Procedure (MRPP), which tests for significant differences between sampling units (method+feature-fixing in this study) [49]. As MRPP compares within and between group dissimilarities, it is similar to multivariate analysis of variance, but does not require data to exhibit multivariate normality [49], making the method appropriate for comparing MSE matrices. MRPP evaluates observed within-group distances relative to average between-group distances of two random groups; the groups may be regarded as significantly different if average within-group distances are less than average between-group distances. MRPP was performed using the *mrpp* function in the R package *vegan* [50]. The analysis returns several values of note: observed delta (δ), the overall mean of group mean distances, weighted by the number of groups; expected delta E(δ), expected delta under the null hypothesis of no group structure; A, a chance-corrected estimate of the proportion of variance explained by group membership; and p, the significance of the test. Each MRPP analysis was run with 999 permutations. The number of permuted between-group distances that are less than the observed within-group δ determines the test’s significance. Additionally, as significant between-group differences may be the result of a difference in means (location) or a difference of variance (dispersion) [51], the homogeneity of variance of each method was also compared. Analysis of multivariate homogeneity of variance was performed with the *betadisper* function in *vegan* [50]; significance was evaluated with Tukey’s Honest Significant Differences.

Due to the positive correlation between MSE and cP distances, methods that reduce cP distances between shapes may reduce the MSE of propagated landmarks. However, this phenomenon is undesirable since reduced cP distances may imply that the method is less sensitive to shape differences. To examine the interaction of MSE and cP distance, we generated matrices of MSE scaled to cP distances (√(MSE)/cP distance), and performed MRPP and homogeneity of variance analyses on this set of matrices as well. Matrix heat maps of MSE, cP distances, and scaled MSE for each method are in the S2 Appendix.

## Results

### Quantifying errors of cPDist

Visual inspection of many pairwise mappings reveals that alignment errors in cPDist occur with undesirable frequency. Of the 583 pairwise mappings inspected, 161 (27.6%) exhibit some type of error. The majority of these errors are relatively subtle and represent some landmark distortion at either the anterior or posterior end of the tooth (trigonid clustering [n=57, 9.8%], talonid drift [41, 7.0%]). The remaining 63 mappings display major errors, either buccal-lingual inversions (33, 5.7%), anterior-posterior inversions (27, 4.6%), or 90° rotations (3, <1%).

Errors generally arose when landmarks were propagated between dissimilar teeth. To our advantage, dissimilarity can be quantified using cP distances. The cP distances between teeth with good maps averaged 0.058 (max=0.116, min=0.025, sd=0.014), while cP distances between teeth with bad maps averaged 0.073 (max=0.114, min=0.046, sd=0.013). The difference between these distributions is highly significant (Tukey’s Q=17.44, *p*<0.001). There are also significant differences between the cP distances of good maps and all particular error types except 90° rotation (Table 2). Although there are no significant differences in the cP distances between any two error classes, the more subtle errors of trigonid clustering and talonid drift tend to occur in pairwise comparisons involving smaller cP distances than the side-to-side inversions or 90° rotation (Fig 2a).

**Fig 2.**
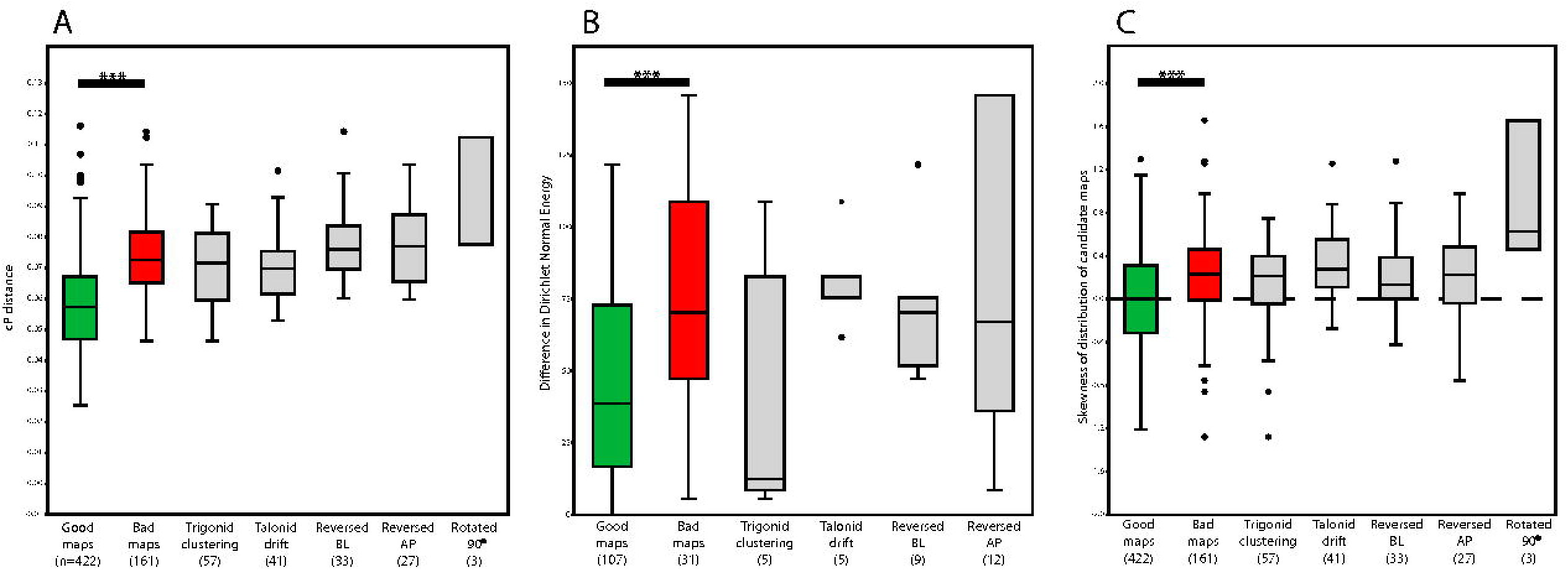
Boxplots of cP distances (A), Dirichlet Normal Energy (B), and skewness of the distribution of candidate maps (C) for good maps, bad maps, and maps of each error type. Asterisks denote significant differences (*p*<0.001) between group means. Boxes include 25-75% quartiles; whiskers extend to furthest points less than 1.5 times the interquartile range. Circles indicate outliers.

**Table 2.**
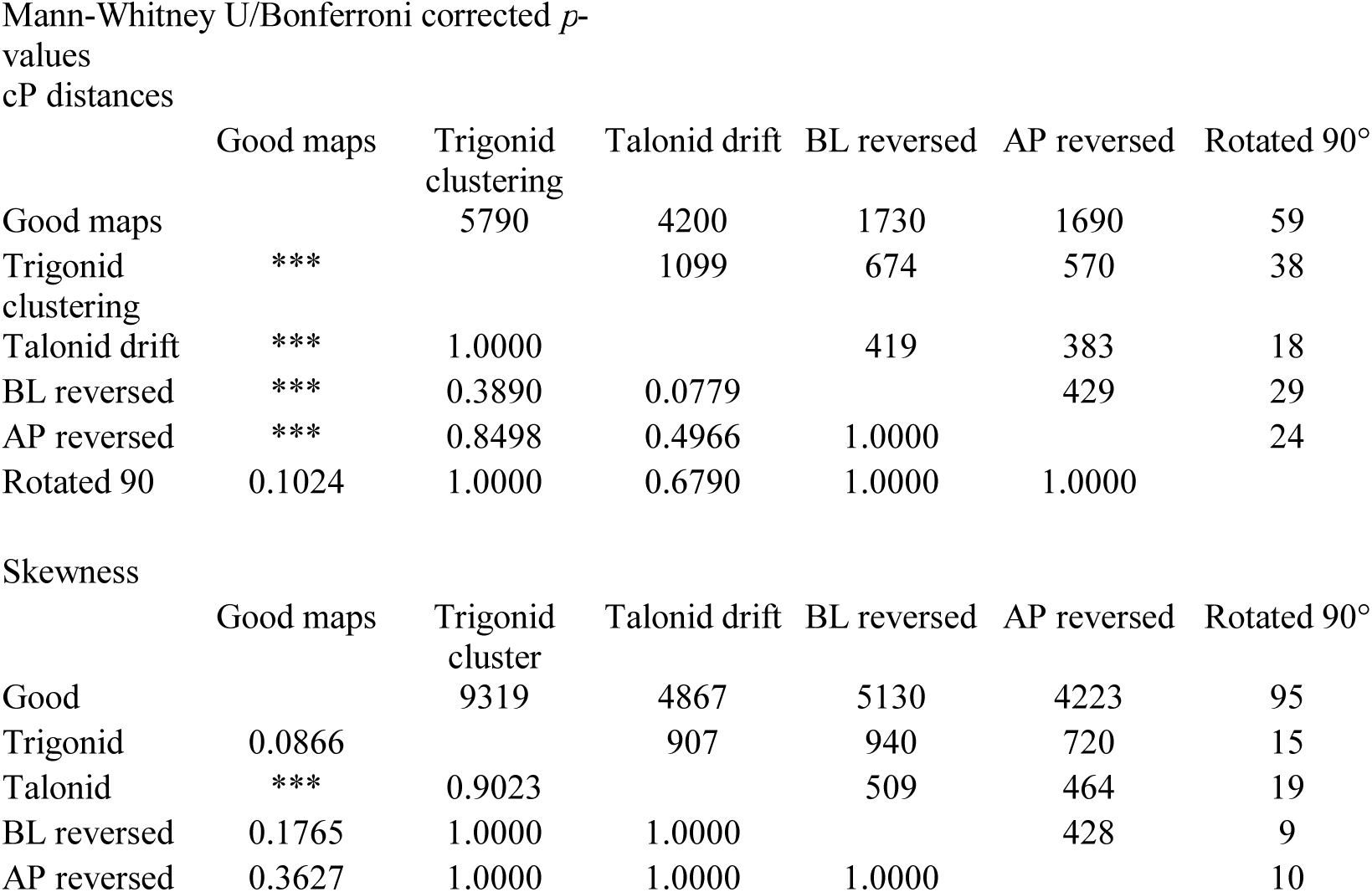
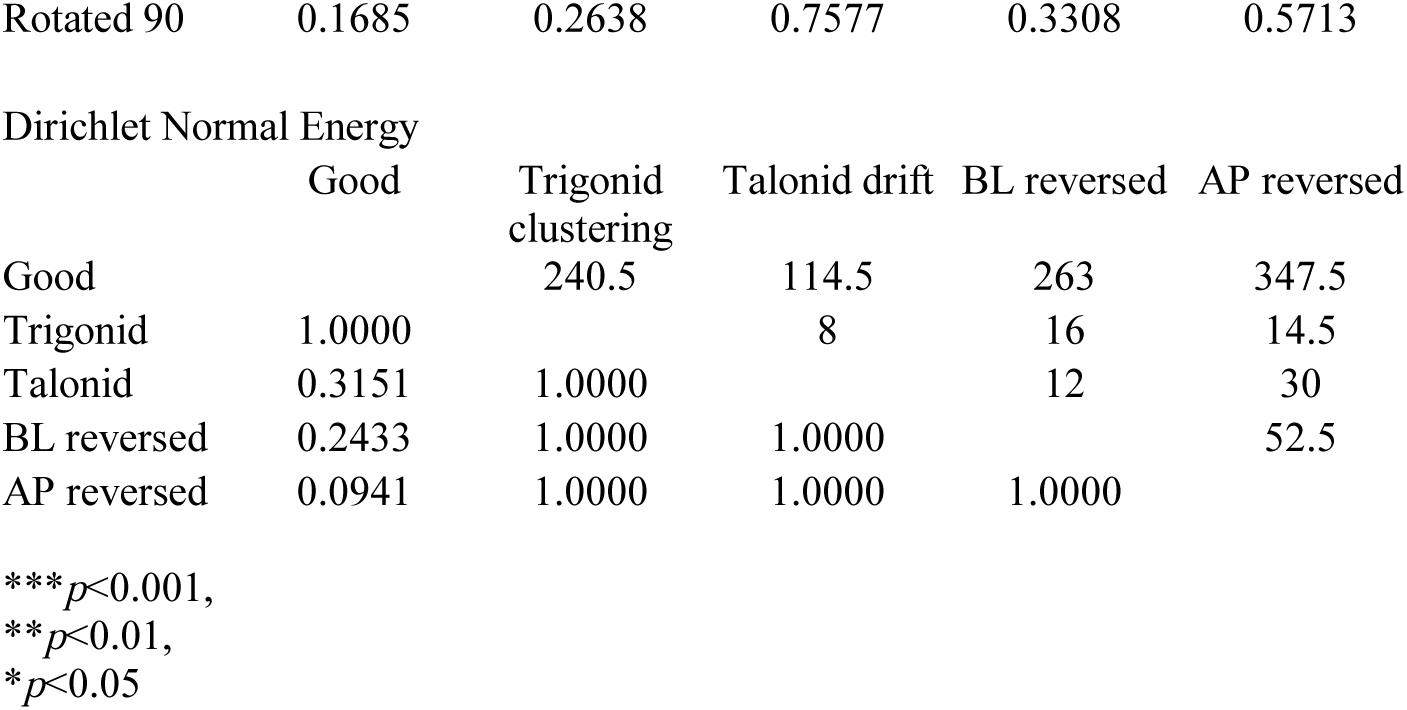
Mann-Whitney U-test results for cP distances, skewness, and Dirichlet Normal Energy (DNE) of good and bad maps.

Mann-Whitney U/Bonferroni corrected $p$ -values cP distances
|  | Good maps | Trigonid clustering | Talonid drift | BL reversed | AP reversed | Rotated 90° |
| --- | --- | --- | --- | --- | --- | --- |
| Good maps |  | 5790 | 4200 | 1730 | 1690 | 59 |
| Trigonid clustering | *** |  | 1099 | 674 | 570 | 38 |
| Talonid drift | *** | 1.0000 |  | 419 | 383 | 18 |
| BL reversed | *** | 0.3890 | 0.0779 |  | 429 | 29 |
| AP reversed | *** | 0.8498 | 0.4966 | 1.0000 |  | 24 |
| Rotated 90 | 0.1024 | 1.0000 | 0.6790 | 1.0000 | 1.0000 |  |

Skewness
|  | Good maps | Trigonid cluster | Talonid drift | BL reversed | AP reversed | Rotated 90° |
| --- | --- | --- | --- | --- | --- | --- |
| Good |  | 9319 | 4867 | 5130 | 4223 | 95 |
| Trigonid | 0.0866 |  | 907 | 940 | 720 | 15 |
| Talonid | *** | 0.9023 |  | 509 | 464 | 19 |
| BL reversed | 0.1765 | 1.0000 | 1.0000 |  | 428 | 9 |
| AP reversed | 0.3627 | 1.0000 | 1.0000 | 1.0000 |  | 10 |

|  |  |  |  |  |  |
| --- | --- | --- | --- | --- | --- |
| Rotated 90 | 0.1685 | 0.2638 | 0.7577 | 0.3308 | 0.5713 |
| Dirichlet Normal Energy |  |  |  |  |  |
|  | Good | Trigonid clustering | Talonid drift | BL reversed | AP reversed |
| Good |  | 240.5 | 114.5 | 263 | 347.5 |
| Trigonid | 1.0000 |  | 8 | 16 | 14.5 |
| Talonid | 0.3151 | 1.0000 |  | 12 | 30 |
| BL reversed | 0.2433 | 1.0000 | 1.0000 |  | 52.5 |
| AP reversed | 0.0941 | 1.0000 | 1.0000 | 1.0000 |  |

\*\*\* $p < 0.001$ ,
\*\* $p < 0.01$ ,
\* $p < 0.05$

The difference in DNE was significantly lower (Tukey’s Q=5.44, *p*<0.001) for good mappings (mean=46.2, max=121.7, min=0, sd=34.5) than for bad maps (mean=74.6, min=5.6, max=145.8, sd=41.5). However, there were no significant differences between DNE contrasts of good mappings and any particular error type (Table 2, Fig 2b). These results confirm our hypothesis that bad mappings occur when landmarks are propagated between highly dissimilar teeth.

Finally, we calculated the skewness of the distributions of the candidate maps for all 583 pairwise mappings. For 422 good mappings, map distributions had a mean skewness of 0.007 (min=-1.21, max=1.30, sd=0.42), which is not significantly different from 0 (*p*=0.84). The 161 bad maps had a mean skewness of 0.22 (min=-1.28, max=1.66, sd=0.39), which is significantly different from 0 (*p*<0.001), indicating that the distribution of candidate maps exhibits significant positive skew when cPDist selects a bad map. Significant differences were recovered in the skewness of good and bad maps (Tukey’s Q=7.88, *p*<0.001), as well as between good maps and those with talonid drift, trigonid clustering, and buccal-lingual inversions (Table 2, Fig 2c).

These results confirm the prediction that good maps are much more distinct from other candidate maps than bad maps. When candidate maps are normally distributed, the map with the smallest cP distance is likely to be an accurate map. When candidate maps are positively skewed, cPDist chooses between several maps with similar cP distances. Given the plurality of possibilities and numerical error, the algorithm is more likely to select a bad map.

### Evaluating accuracy of a minimum spanning tree approach

Visual inspection of propagated landmark errors reveals that bad maps are generated when two shapes are quite dissimilar from one another. If any of the branches of the MST utilized by cPMST connect dissimilar shapes, the global map could also do a poor job propagating landmarks. Given the distribution of cP distances of all 161 observed bad maps, cP distances of less than 0.047 are outside the 95% confidence interval (mean=0.073, sd=0.013). Only one inspected bad map had a cP distance less than 0.047; this map exhibited trigonid clustering, a relatively minor propagation error. Of the 115 branches in the MST, only 3 branches (2.6%) had cP distances greater than 0.047. Additionally, only one MST branch had a cP distance greater than the mean of all observed good mappings (0.062, *Galago senegalensis* K05 – *Cynocephalus volans* u16). Thus, it seems likely that propagating landmarks through the MST greatly reduces the likelihood of serious errors such as inversions or rotations. Fig 3 compares the landmark MSEs of cPDist with cPMST (without feature-fixing), and demonstrates that the latter has a much lower MSE than the former. However, as discussed above, the MST approach also has the undesirable property of increasing landmark MSE between shapes that are similar; we evaluate the accuracy of the novel globally informed methods relative to cPDist below.

**Fig 3.**
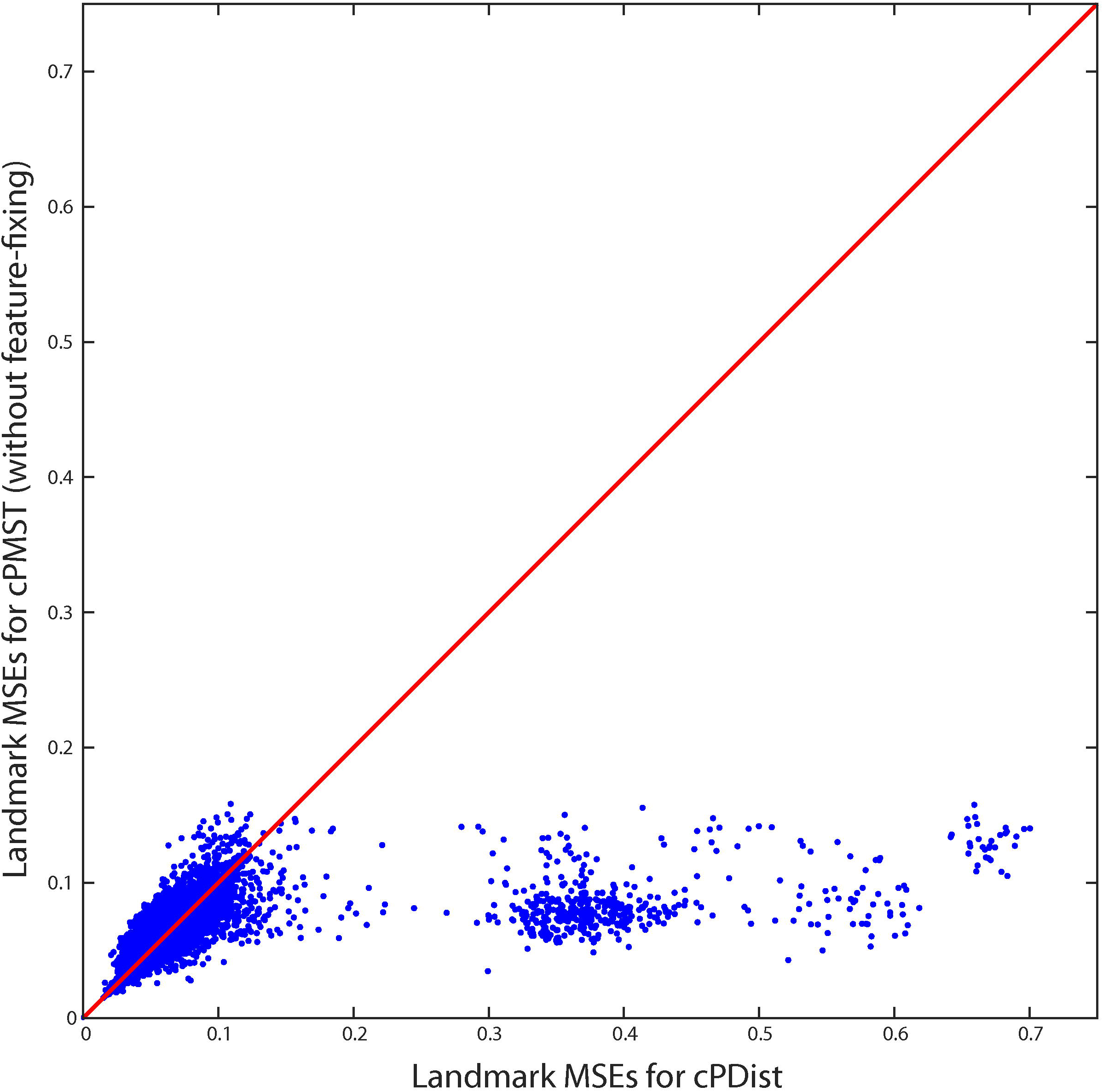
Pairwise comparisons of landmark MSE generated by cPDist and cPMST (without feature-fixing). For many pairwise comparisons, cPMST (without feature-fixing) has much lower landmark MSE than cPDist. In particular, large (>0.1583) MSEs are reduced with cPMST (without feature-fixing) compare to cPDist. Red line indicates line of equivalence (y=x).

### Comparing effects of globally informed methods on the characterization of geometric affinities

Of the five one-way ANOVAs run (Table 3), only sampling resolution was non-significant. The most significant factor was feature-fixing (*p*<<0.0001). The method of sequential comparison (MST v. LAST or Viterbi, etc.) was also highly significant (*p*=0.0005), followed by the propagation root of pseudolandmarks (*p*=0.005), and composedness for LAST trees (*p*=0.01).

**Table 3.** Results of one-way ANOVA assessing parameter effects on shape space.

| Source | Sum Sq. | DF | Mean Sum Sq. | F-value | p-value |
| --- | --- | --- | --- | --- | --- |
| Groups | 9.0277 | 7 | 1.2897 | 4.0731 | *** |
| Error | 35.4626 | 112 | 0.3166 |  |  |
| Total | 44.4902 | 119 |  |  |  |

Feature-Fix
| Source | Sum Sq. | DF | Mean Sum Sq. | F-value | p-value |
| --- | --- | --- | --- | --- | --- |
| Groups | 6.9913 | 1 | 6.9913 | 21.9998 | *** |
| Error | 37.499 | 118 | 0.3178 |  |  |
| Total | 44.4902 | 119 |  |  |  |

Root
| Source | Sum Sq. | DF | Mean Sum Sq. | F-value | p-value |
| --- | --- | --- | --- | --- | --- |
| Groups | 2.867 | 1 | 2.867 | 8.1278 | ** |
| Error | 41.6232 | 118 | 0.3527 |  |  |
| Total | 44.4902 | 119 |  |  |  |

Resolution
| Source | Sum Sq. | DF | Mean Sum Sq. | F-value | p-value |
| --- | --- | --- | --- | --- | --- |
| Groups | 0.1671 | 2 | 0.0836 | 0.22057 | 0.80239 |
| Error | 44.3231 | 117 | 0.3788 |  |  |
| Total | 44.4902 | 119 |  |  |  |

Composedness (48 methods only)
| Source | Sum Sq. | DF | Mean Sum Sq. | F-value | p-value |
| --- | --- | --- | --- | --- | --- |
| Groups | 2.1184 | 1 | 2.1184 | 7.097 | * |
| Error | 13.7304 | 46 | 0.2985 |  |  |
| Total | 15.8487 | 47 |  |  |  |

\*\*\* $p < 0.001$ , \*\* $p < 0.01$ , \* $p < 0.05$

Two-way ANOVAs were used to determine whether interaction effects existed between any pairs of factors. Because our assessment of error in propagated user-determined landmarks was explained largely by the choice of different methods (see results of linear mixed model analysis below), we checked for interaction effects between method and each of the other factors. Significant interaction effects were found between method and feature-fixing, as well as method and composedness (Table 4). Neither propagation root nor sampling resolution interacted significantly with method type (Table 4). In order to include composedness in the two-way ANOVA with method, we were limited to using only the 48 data points that included balanced distribution of method types for each of the composed groups.

**Table 4.** Results of two-way ANOVA assessing parameter effects on shape space.

Method vs. Feature-fix
| Source | Sum Sq. | DF | Mean Sum Sq. | F-value | p-value |
| --- | --- | --- | --- | --- | --- |
| Method | 9.0277 | 7 | 1.2897 | 5.8952 | *** |
| Feature-fix | 6.9913 | 1 | 6.9913 | 31.9575 | *** |
| Method:Feature-fix | 5.7195 | 7 | 0.8171 | 3.7349 | *** |
| Error | 22.7518 | 104 | 0.2188 |  |  |
| Total | 44.4902 | 119 |  |  |  |

Method vs. Root
| Source | Sum Sq. | DF | Mean Sum Sq. | F-value | p-value |
| --- | --- | --- | --- | --- | --- |
| Method | 9.0277 | 7 | 1.2897 | 4.6674 | *** |
| Root | 2.8670 | 1 | 2.8670 | 10.3758 | ** |
| Method:Root | 3.8587 | 7 | 0.5512 | 1.9949 | 0.0627 |
| Error | 28.7369 | 104 | 0.2763 |  |  |
| Total | 44.4902 | 119 |  |  |  |

Method vs. Resolution
| Source | Sum Sq. | DF | Mean Sum Sq. | F-value | p-value |
| --- | --- | --- | --- | --- | --- |
| Method | 9.0277 | 7 | 1.2897 | 3.5524 | ** |
| Resolution | 0.1671 | 2 | 0.0836 | 0.2302 | 0.7948 |
| Method:Resolution | 0.4435 | 14 | 0.0317 | 0.0873 | 1.0000 |
| Error | 34.8519 | 96 | 0.3630 |  |  |
| Total | 44.4902 | 119 |  |  |  |

Method vs. Composedness (only 48 methods)
| Source | Sum Sq. | DF | Mean Sum Sq. | F-value | p-value |
| --- | --- | --- | --- | --- | --- |
| method | 1.5290 | 1 | 1.5290 | 6.3377 | * |
| composedness | 2.1184 | 1 | 2.1184 | 8.7806 | ** |
| method:composedness | 1.5862 | 1 | 1.5862 | 6.5746 | * |
| Error | 10.6152 | 44 | 0.2413 |  |  |
| Total | 15.8487 | 47 |  |  |  |

\*\*\*p<0.001,\*\*p<0.01,\*p<0.05

A linear mixed model including all factors and interaction effects (Table 5) shows a pattern of relative significance comparable to ANOVA results, although method explains more variance than feature-fixing does. Sequentially dropping non-significant terms and re-running the analysis leads to a final model with three terms, included method, feature-fixing, and an interaction term between the two (Table 5). This result is strongly consistent with ANOVA results (Table 3-4).

**Table 5.** Linear mixed model for assessment of parameters explain variance from user-based landmark approach. The initial model included all four factors (method, feature-fixing, root, and resolution) and interactions between method and the other three factors. Non-significant factors were sequentially dropped to arrive at the final model.

| Factor | Sum Sq. | DF | Mean Sum Sq. | F-value | p-value |
| --- | --- | --- | --- | --- | --- |
| Method | 0.4086 | 7 | 0.0584 | 4.7011 | *** |
| Feature-fix | 0.0966 | 1 | 0.0966 | 7.7771 | ** |
| Method:Feature-fix | 0.2452 | 7 | 0.0350 | 2.8211 | * |
| Root | 0.0402 | 1 | 0.0402 | 3.2365 | 0.0758 |
| Resolution | 0.0349 | 2 | 0.0174 | 1.4040 | 0.2516 |
| Method:Root | 0.0704 | 7 | 0.0101 | 0.8102 | 0.5814 |
| Method:Resolution | 0.0541 | 14 | 0.0039 | 0.3113 | 0.9911 |
| Error | 0.9934 | 80 | 0.0124 |  |  |

Final model
| Factor | Sum Sq. | DF | Mean Sum Sq. | F-value | p-value |
| --- | --- | --- | --- | --- | --- |
| Method | 0.4086 | 7 | 0.0584 | 5.0890 | *** |
| Feature-fix | 0.0966 | 1 | 0.0966 | 8.4187 | ** |
| Method:Feature-fix | 0.2452 | 7 | 0.0350 | 3.0538 | ** |
| Error | 1.1930 | 104 | 0.0115 |  |  |

\*\*\*p<0.001,\*\*p<0.01,\*p<0.05

The ANOVA and linear mixed model results highlight the importance of method, feature-fixing, and their interaction. To further assess how feature-fixing influence method effects, we split the dataset into two subsets. The first subset contained analyses that used feature-fixing, while the second subset contained analyses that did *not* use feature-fixing. We then ran one-way ANOVA with method as the factor. Results of this one-way ANOVA indicate that method is much more significant when feature-fixing is not used (Table 6).

**Table 6.**
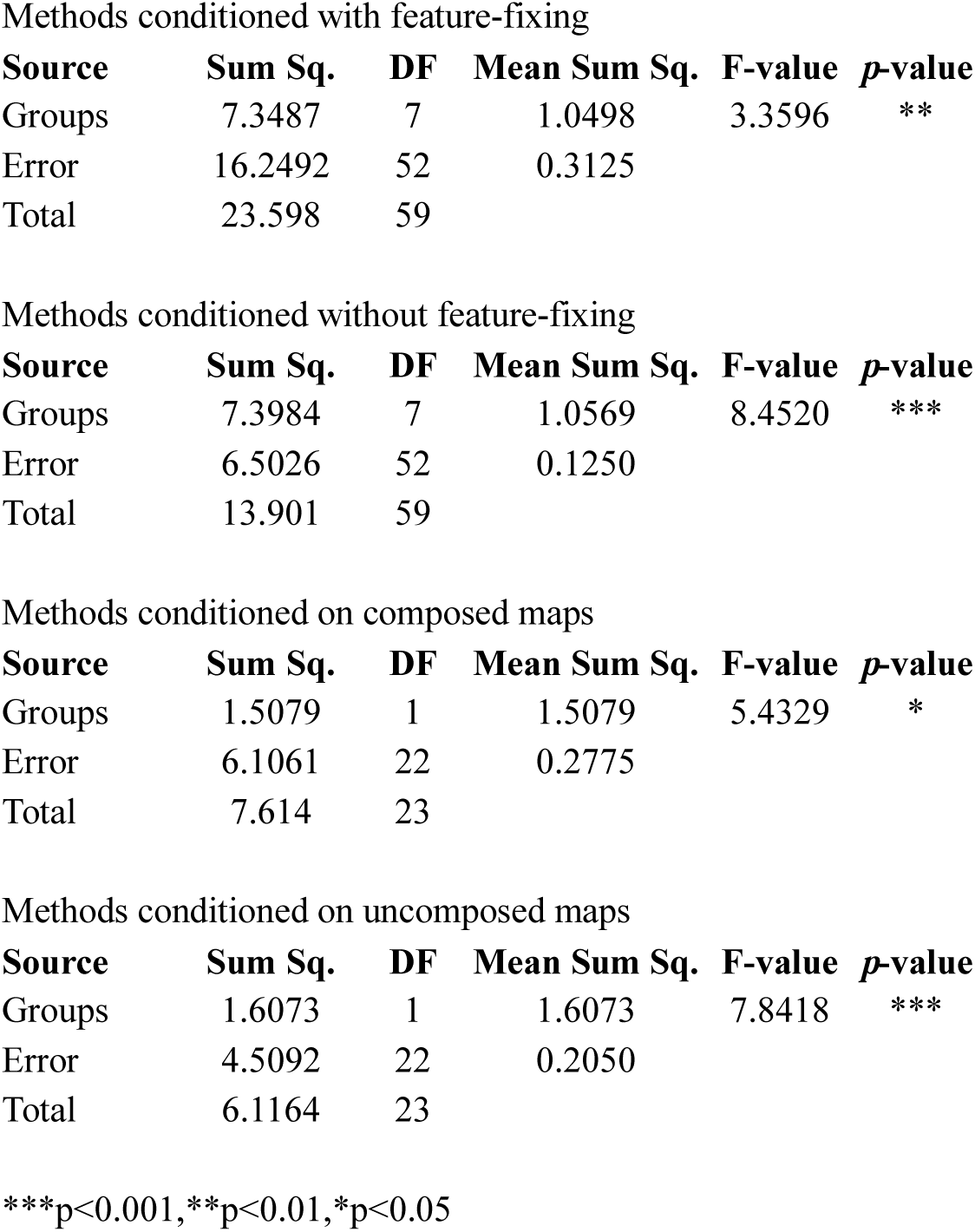
Conditional ANOVAs for assessment of interaction effects between method and feature-fixing, and method and composedness.

Due to unequal sample sizes, we could not test the effect of composedness in our linear model. However, composedness was a significant parameter in one-way ANOVA and exhibited significant interaction effects with method. Splitting the dataset into composed and uncomposed subsets and running one-way ANOVAs with method as a factor indicates that method is more significant on uncomposed maps (Table 6).

### Evaluating accuracy of globally informed methods

With the exceptions of LAST (α = balance) and the Viterbi methods (with or without feature-fixing), all of the proposed globally informed methods have much lower maximum landmark MSEs than cPDist, suggesting that these globally informed methods successfully avoid large-scale misalignments and correspondingly elevated landmark MSE between user-determined and propogated landmarks (Table 7). Since LAST (α = balance) and the Viterbi methods (with or without feature-fixing) have maximum landmark MSEs that are comparable to the maximum landmark MSE of cPDist, these methods may be susceptible to misalignments of similar magnitude as cPDist. Among the novel methods with much lower maximum landmark MSEs than cPDist, only composed LAST (α = median; no feature-fixing) has a lower maximum landmark MSE value than the performance baseline established by cPMST (without feature-fixing) (maximum landmark MSEs = 0.1507 and 0.1583 respectively). Still, many other methods that do not utilize feature-fixing (e.g., LAST [α = mean], composed LAST [α = balance], composed LAST [α = mean]) have similarly low maximum landmark MSEs (Table 7).

**Table 7.** Summary statistics of differences in landmark MSE under each globally informed method relative to cPDist. Positive residuals indicate the globally informed method has greater landmark MSE than cPDist, while negative residuals indicate globally informed method has smaller landmark MSE than cPDist. “Maximum landmark MSE” indicates the inflection point above which the globally informed method always decreases MSE relative to cPDist. Note that the % of points above or below the line y=x may not sum to 100, as some points may be on the line. Bold text represents maximum or minimum values as appropriate. SD, standard deviation.

| Method | Variant | Feature-fixing | Maximum landmark MSE | % of points above $y=x$ | Positive residuals | | | | % of points below $y=x$ | Negative residuals | | | |
| --- | --- | --- | --- | --- | --- | --- | --- | --- | --- | --- | --- | --- | --- |
|  |  |  |  |  | Mean | Median | Maximum | SD |  | Mean | Median | Maximum | SD |
| cPDist | - | N | 0.7003 | na | na | na | na | na | na | na | na | na | na |
| cPMST | - | N | 0.1583 | 56.0 | **0.0092** | **0.0070** | 0.0645 | 0.0082 | 42.6 | 0.0629 | 0.0082 | 0.5785 | 0.1239 |
| LAST | Mean | N | 0.1617 | 55.2 | 0.0115 | 0.0082 | 0.0787 | 0.0106 | 43.4 | 0.0623 | 0.0086 | 0.5785 | 0.1230 |
| LAST | Balance | N | 0.6901 | 55.2 | 0.0170 | 0.0059 | 0.6048 | 0.0641 | 43.3 | 0.0526 | 0.0060 | 0.5500 | 0.1099 |
| Composed LAST | 1 | N | 0.4488 | **66.3** | 0.0313 | 0.0106 | 0.3844 | 0.0721 | **32.4** | 0.0628 | 0.0093 | 0.6033 | 0.1272 |
| Composed LAST | Balance | N | 0.1583 | 56.0 | **0.0092** | **0.0070** | 0.0645 | 0.0082 | 42.6 | 0.0629 | 0.0082 | 0.5785 | 0.1239 |
| Composed LAST | Mean | N | 0.1627 | 65.3 | 0.0164 | 0.0145 | 0.0652 | 0.0123 | 33.3 | **0.0773** | **0.0103** | 0.5785 | **0.1350** |
| Composed LAST | Median | N | **0.1507** | 57.6 | 0.0098 | 0.0078 | **0.0573** | **0.0080** | 41.0 | 0.0653 | 0.0085 | 0.5785 | 0.1261 |
| Viterbi | 0.0 | N | 0.6801 | 35.1 | 0.0169 | 0.0066 | 0.4930 | 0.0455 | 41.8 | 0.0479 | 0.0072 | 0.5972 | 0.1101 |
| Viterbi | 0.25 | N | 0.6679 | 35.4 | 0.0131 | 0.0060 | 0.4658 | 0.0366 | 42.9 | 0.0461 | 0.0072 | 0.6026 | 0.1071 |
| Viterbi | 0.5 | N | 0.6801 | 35.9 | 0.0131 | 0.0060 | 0.4658 | 0.0362 | 42.7 | 0.0461 | 0.0071 | **0.6165** | 0.1080 |
| cPMST | - | Y | 0.1854 | 45.8 | 0.0099 | 0.0076 | 0.0752 | 0.0088 | 54.2 | 0.0528 | 0.0096 | 0.5886 | 0.1125 |
| LAST | Mean | Y | 0.1968 | 47.9 | 0.0109 | 0.0082 | 0.0810 | 0.0100 | 52.1 | 0.0548 | 0.0099 | 0.5886 | 0.1141 |
| LAST | Balance | Y | 0.6737 | 43.5 | 0.0192 | 0.0072 | 0.5869 | 0.0655 | 56.5 | 0.0443 | 0.0091 | 0.5464 | 0.0975 |
| Composed LAST | 1 | Y | 0.4750 | 55.5 | 0.0341 | 0.0102 | 0.3825 | 0.0762 | 44.5 | 0.0488 | 0.0091 | 0.6085 | 0.1110 |
| Composed LAST | Balance | Y | 0.1854 | 45.8 | 0.0099 | 0.0076 | 0.0752 | 0.0088 | 54.2 | 0.0528 | 0.0096 | 0.5886 | 0.1125 |
| Composed LAST | Mean | Y | 0.1887 | 52.1 | 0.0118 | 0.0092 | 0.0726 | 0.0102 | 47.9 | 0.0578 | 0.0101 | 0.5967 | 0.1174 |
| Composed LAST | Median | Y | 0.1796 | 48.3 | 0.0105 | 0.0083 | 0.0809 | 0.0091 | 51.7 | 0.0546 | 0.0095 | 0.5886 | 0.1146 |
| Viterbi | 0.0 | Y | 0.6844 | 44.2 | 0.0147 | 0.0068 | 0.4872 | 0.0396 | 55.8 | 0.0385 | 0.0087 | 0.6129 | 0.0964 |
| Viterbi | 0.25 | Y | 0.6163 | 43.5 | 0.0124 | 0.0066 | 0.4876 | 0.0325 | 56.5 | 0.0378 | 0.0088 | 0.5949 | 0.0944 |
| Viterbi | 0.5 | Y | 0.6844 | 43.7 | 0.0124 | 0.0065 | 0.4876 | 0.0321 | 56.3 | 0.0378 | 0.0087 | 0.6117 | 0.0950 |

Two trends are apparent when feature-fixing is implemented: 1) methods with feature-fixing have greater maximum landmark MSE than the same method without feature-fixing, and 2) methods with feature-fixing are appear to be less susceptible to the accumulation of numerical error during landmark propagation, as indicated by the percentage of points above the line y=x (Table 7). These positive residuals indicate pairwise mappings in which landmark MSE has increased for the novel method relative to the landmark MSE observed for cPDist (Fig 3 provides an example for cPMST). While no single method has the lowest mean, median, or maximum positive residual, the set of cPMST, LAST (α = mean), and three composed LAST (α = balance, mean, or median) methods share low positive residuals with low dispersion (Table 7). When feature-fixing is implemented with these methods, both the magnitude and dispersion of these residuals increase, but the number of positive residuals decreases. The decrease in the number of positive residuals is observed for all methods except the Viterbi methods.

Compared to cPMST, Viterbi methods (with or without feature-fixing) have smaller median positive residuals and larger mean positive residuals. Based on the percentage of points above y=x, Viterbi methods (with or without feature-fixing) experience less inflation of landmark MSE during landmark propagation (Table 7). However, all Viterbi methods have large maximum positive residuals and high variance in the distribution of these residuals. Further, while the Viterbi methods have the greatest number of points below y=x (indicating that cPDist has greater landmark MSE), the means and medians of these negative residuals are smaller and the standard deviations are lower than the values recovered for cPMST. So while Viterbi methods experience less inflation of landmark MSE due to the accumulation of numerical error, these methods are likely to be more prone to large-scale misalignments (similar to cPDist). Essentially, compared to all other methods, landmark MSEs of the Viterbi methods have higher correlations with the landmark MSE of cPDist, which can be seen in bivariate plots of the landmark MSEs of these methods (S2 Appendix).

Results from multivariate homogeneity of variance tests confirm significant differences in the dispersion of landmark MSE under different methods (S1 Appendix, S2 Appendix). cPDist has the highest variance (measured as the Euclidean distance from each matrix entry to the centroid of the matrix), and all Viterbi methods exhibit similarly high dispersion. The variance of landmark MSEs for cPDist and all Viterbi methods is significantly greater than any other method, and cPDist has significantly greater variance than all Viterbi methods (S1 Appendix, S2 Appendix). A slightly different pattern of variance emerges when MSEs are scaled to pairwise cP distances (S1 Appendix, S2 Appendix). cPDist maintains the greatest dispersion, but all methods that utilize feature-fixing have reduced variance relative to their non-feature-fixing counterparts (S1 Appendix, S2 Appendix). In most cases, pairwise comparisons of the same method with and without feature-fixing produce significant differences in terms of variance; methods with feature-fixing always have lower variance than methods without. Viterbi methods without feature-fixing also have relatively high dispersion, similar to the unscaled MSE results.

The observed heterogeneity of variance renders any test of significant differences in method means suspect. With this caveat, MRPP indicates there are significant differences between methods, as gauged by both MSE and scaled MSE comparisons (MSE: δ=0.47, E(δ)=0.60, A=0.21, *p*=0.001; scaled MSE: δ=12.8, E(δ)=14.64, A=0.13, *p*=0.001). Both within-and between-method MSE and scaled MSE are detailed in the S1 Appendix. To compensate for the observed heterogeneity of variance, we also performed MRPP analysis for a reduced sample of MSE matrices, excluding all methods with significantly higher dispersion (cPDist, all six Viterbi methods). In this analysis, significant differences between methods were still recovered (δ=0.31, E(δ)=0.47, A=0.34, *p*=0.001). The observed patterns of heterogeneity in scaled MSE comparisons were too diffuse to permit a similarly restricted analysis.

Table 8 provides summary statistics for cP distances, landmark MSE, and scaled MSE for all methods. cPMST (with feature-fixing) has the lowest mean landmark MSE (0.062), while LAST (α = mean, feature-fixing) has the highest mean cP distance and the lowest mean scaled MSE (3.165). When considering only those methods without feature-fixing, the set of cPMST, LAST (α = mean), and three composed LAST (α = balance, mean, or median) methods share similar characteristics: moderate mean and maximum cP distances, low mean and maximum landmark MSEs, and the lowest mean and maximum scaled MSEs (Table 8). Compared to other methods without feature-fixing, this set of methods also exhibits lower standard deviations for landmark and scaled MSE.

**Table 8.**
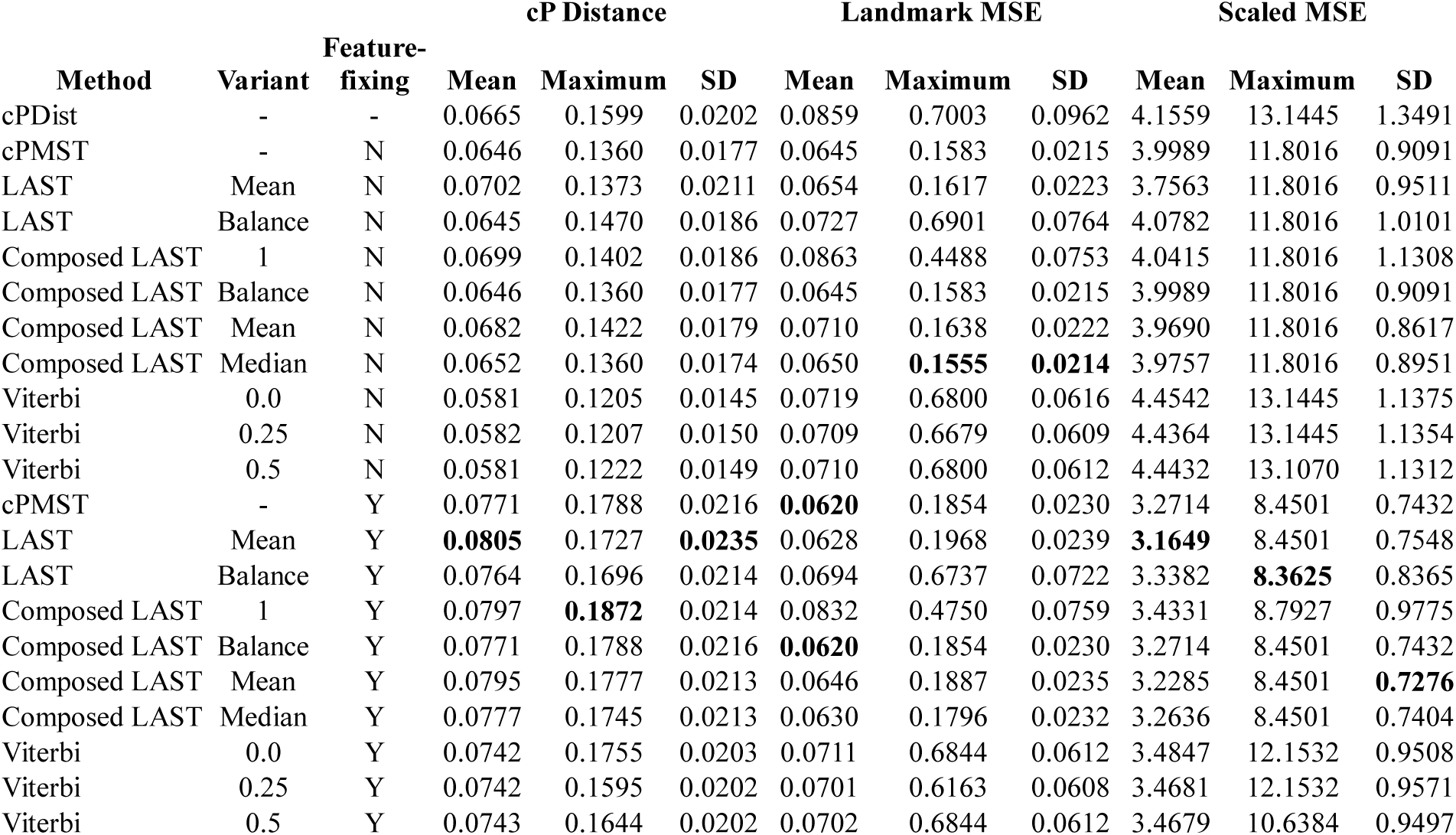
Summary statistics of pairwise cPdistances, landmark MSE, and scaled MSE by method. Bold text represents maximum or minimum values as appropriate. MSE, mean square error; SD, standard deviation.

| Method | Variant | Feature-fixing | cP Distance |  |  | Landmark MSE |  |  | Scaled MSE |  |  |
| --- | --- | --- | --- | --- | --- | --- | --- | --- | --- | --- | --- |
|  |  |  | Mean | Maximum | SD | Mean | Maximum | SD | Mean | Maximum | SD |
| cPDist | - | - | 0.0665 | 0.1599 | 0.0202 | 0.0859 | 0.7003 | 0.0962 | 4.1559 | 13.1445 | 1.3491 |
| cPMST | - | N | 0.0646 | 0.1360 | 0.0177 | 0.0645 | 0.1583 | 0.0215 | 3.9989 | 11.8016 | 0.9091 |
| LAST | Mean | N | 0.0702 | 0.1373 | 0.0211 | 0.0654 | 0.1617 | 0.0223 | 3.7563 | 11.8016 | 0.9511 |
| LAST | Balance | N | 0.0645 | 0.1470 | 0.0186 | 0.0727 | 0.6901 | 0.0764 | 4.0782 | 11.8016 | 1.0101 |
| Composed LAST | 1 | N | 0.0699 | 0.1402 | 0.0186 | 0.0863 | 0.4488 | 0.0753 | 4.0415 | 11.8016 | 1.1308 |
| Composed LAST | Balance | N | 0.0646 | 0.1360 | 0.0177 | 0.0645 | 0.1583 | 0.0215 | 3.9989 | 11.8016 | 0.9091 |
| Composed LAST | Mean | N | 0.0682 | 0.1422 | 0.0179 | 0.0710 | 0.1638 | 0.0222 | 3.9690 | 11.8016 | 0.8617 |
| Composed LAST | Median | N | 0.0652 | 0.1360 | 0.0174 | 0.0650 | **0.1555** | **0.0214** | 3.9757 | 11.8016 | 0.8951 |
| Viterbi | 0.0 | N | 0.0581 | 0.1205 | 0.0145 | 0.0719 | 0.6800 | 0.0616 | 4.4542 | 13.1445 | 1.1375 |
| Viterbi | 0.25 | N | 0.0582 | 0.1207 | 0.0150 | 0.0709 | 0.6679 | 0.0609 | 4.4364 | 13.1445 | 1.1354 |
| Viterbi | 0.5 | N | 0.0581 | 0.1222 | 0.0149 | 0.0710 | 0.6800 | 0.0612 | 4.4432 | 13.1070 | 1.1312 |
| cPMST | - | Y | 0.0771 | 0.1788 | 0.0216 | **0.0620** | 0.1854 | 0.0230 | 3.2714 | 8.4501 | 0.7432 |
| LAST | Mean | Y | **0.0805** | 0.1727 | **0.0235** | 0.0628 | 0.1968 | 0.0239 | **3.1649** | 8.4501 | 0.7548 |
| LAST | Balance | Y | 0.0764 | 0.1696 | 0.0214 | 0.0694 | 0.6737 | 0.0722 | 3.3382 | **8.3625** | 0.8365 |
| Composed LAST | 1 | Y | 0.0797 | **0.1872** | 0.0214 | 0.0832 | 0.4750 | 0.0759 | 3.4331 | 8.7927 | 0.9775 |
| Composed LAST | Balance | Y | 0.0771 | 0.1788 | 0.0216 | **0.0620** | 0.1854 | 0.0230 | 3.2714 | 8.4501 | 0.7432 |
| Composed LAST | Mean | Y | 0.0795 | 0.1777 | 0.0213 | 0.0646 | 0.1887 | 0.0235 | 3.2285 | 8.4501 | **0.7276** |
| Composed LAST | Median | Y | 0.0777 | 0.1745 | 0.0213 | 0.0630 | 0.1796 | 0.0232 | 3.2636 | 8.4501 | 0.7404 |
| Viterbi | 0.0 | Y | 0.0742 | 0.1755 | 0.0203 | 0.0711 | 0.6844 | 0.0612 | 3.4847 | 12.1532 | 0.9508 |
| Viterbi | 0.25 | Y | 0.0742 | 0.1595 | 0.0202 | 0.0701 | 0.6163 | 0.0608 | 3.4681 | 12.1532 | 0.9571 |
| Viterbi | 0.5 | Y | 0.0743 | 0.1644 | 0.0202 | 0.0702 | 0.6844 | 0.0612 | 3.4679 | 10.6384 | 0.9497 |

There are several trends across methods when feature-fixing is implemented. Relative to the same method without feature-fixing, the mean, maximum, and standard deviation of cP distances increase for all methods with feature-fixing (Table 8). For scaled MSE, the mean, maximum, and standard deviation all decrease with feature-fixing. For landmark MSE, not all methods exhibit the same trends when feature-fixing is implemented. While mean landmark MSE decreases with feature-fixing for all methods, maximum landmark MSE increases for all methods except LAST (α = balance) and Viterbi (angle weight = 25%). The variance of landmark MSE increases for all methods except LAST (α = balance) and all three Viterbi methods (Table 8). Finally, while the Viterbi methods (with or without feature-fixing) have the lowest mean scaled MSE, they share the highest maximum scaled MSE with cPDist, and have high variance relative to other methods (Table 8).

## Discussion

### Quantifying errors of cPDist

When comparing highly dissimilar shapes, the pairwise correspondence found by cPDist is particularly error-prone in terms of landmark MSE. This substantial error rate presents a minor paradox: despite the errors lurking in the dataset, Boyer et al. [1] still recovered a strong correlation between user-determined and automatically determined distances and had a nearly equivalent success rate in taxonomic classification. We reconcile this paradox by noting that errors of cPDist occur most often between geometrically dissimilar shapes, resulting in large cP distances due to the restricted search space (conformal maps plus TPS) as opposed to faithfully capturing the geometric dissimilarity. Consequently, cPDist gets the “right answer” for the wrong reasons when comparing very different shapes. Furthermore, the classification method used in [1] assessed only whether small cP distances are taxonomically reliable. Since we have confirmed that only large cP distances are unreliable, there is no real contradiction between our results and Boyer et al.’s [1] assessment. However, it highlights that the approach of [1] does not provide sufficiently informative pairwise correspondences, which are essential for geometric morphological studies.

### Evaluating accuracy of a minimum spanning tree approach

Since all edges of the MST have smaller cP distances than the minimum cP distance of the observed bad maps, it is quite likely that using a MST sufficiently addresses the issue of aligning structures with very different morphologies. In our test case, branches within the MST were small enough to alleviate concerns of major misalignments. However, more work should be done on a wider variety of data sets to determine more precisely the sample properties (e.g., the morphological gaps between objects or the range of morphologies) that increase risk for misalignments even with MST. Additionally, cPMST has the unfortunate side effect of increasing landmark propagation error (reflected in the cP distances) when two similar shapes are not directly connected by an edge of the MST. This issue is partially addressed by subsequent tree-based methods and is discussed further below.

### Comparing effects of globally informed methods on the characterization of geometric affinities

Using ANOVA and linear mixed models, we ascertained the effects of five different factors on the characterization of shape affinities by the globally informed approaches developed for this study (Tables 3-5). The strongest effects were produced by the choice of method and whether or not feature-fixing was used. These two factors had an interaction effect such that including feature-fixing in the protocol reduced the impact of method. From this information alone, it is unclear if feature-fixing is beneficial. Reduction of the method effect through feature-fixing may be beneficial if the results became both more consistent and biologically meaningful. On the other hand, feature-fixing may increase random error in the results, creating the observed statistical effect, thus leading to more variable and less biologically meaningful results. To determine which is more likely for this particular dataset, we assess shape space characterization of a few example methods below.

The root for pseudolandmark propagation had little obvious effect on the results (Tables 3-5). Our analyses suggest that the best approach is to use the specimen with the minimum average difference from all other specimens in the collection as the starting point for landmark propagation. Thought the root shape chosen this way is *not* the same among all 120 analyses, no significant interaction between the root and the choice of method was recovered (Tables 4-5). If root had a strong direct effect, we should observe a significant interaction between root and method (since the root was different under each method). Though the effect of the root was minimal in this analysis, it may still be an important parameter for other datasets, particularly those that include highly dissimilar shapes (e.g., the combined astragalus and calcaneus dataset of [2]).

For our dataset, the impact of composed vs. uncomposed maps seems more important than the root, since the composedness has a significant interaction with method. However, as stated above, it could not be included in the linear model due to unequal sample sizes (and unbalanced experiment design) relative to other factors. One-way ANOVA run on method using a composed/uncomposed split sample suggests that composed maps tend to decrease differences between methods. Therefore, it is probably more desirable to account for cP distances measured for composed maps when sequentially concatenating pairwise correspondences between similar shapes (unless there is a reason to prefer properties of an individual method).

Finally, pseudolandmark sampling density had virtually no effect. This is highly encouraging, and suggests that relatively fewer pseudolandmarks may be used to decrease the computational intensity of downstream analyses.

### Evaluating accuracy of globally informed methods

The alternative tree-based approaches presented here were developed primarily to minimize the accumulation of numerical error as landmarks are propagated through a tree. Based on the analysis of landmark MSE residuals above, a set of methods perform similar to cPMST, including LAST (α = mean) and three composed LAST (α = balance, mean, or median) methods. While none of these approaches has substantially better performance than cPMST, composed LAST (α = median) does exhibit fewer positive landmark MSE residuals, a lower maximum positive residual, and lower variance of these residuals (Table 7), indicating that it may be somewhat preferable to cPMST. In addition, with or without feature-fixing, composed LAST (α = median) exhibits a higher mean cP distance and lower scaled MSE than cPMST (Table 8).

Because feature-fixing aims to match geometric characteristics on the shapes (e.g. cusp tips or basins) and many user-determined landmarks lie near these positions, we expected feature-fixing to reduce landmark MSE. However, based on the analysis of MSE residuals, feature-fixing appears to have a mixed effect on landmark MSE. The procedure tends to increase maximum landmark MSE, but decrease the number of positive MSE residuals (for all except the Viterbi methods) (Table 7). Further, because feature-fixing increases cP distances, scaled MSEs are lower when feature-fixing is implemented (Table 8). Thus, based on landmark MSE, the potential benefits of feature-fixing are ambiguous.

Multivariate homogeneity of variance tests performed on MSE matrices (S1 Appendix) reveal that cPDist and all six Viterbi methods have significantly greater variance than any other method. This result is not surprising, as visual inspection of pairwise cPDist mappings revealed multiple instances of propagation error (reflected by high landmark MSEs). Reduced variance in the MST-based methods suggests that erroneous mappings have been largely eliminated, which was a primary goal in developing these subsequent tree-based methods. The Viterbi methods are different from other tree-based methods since they aim to minimize the distance functional but sacrifice global transitivity.

Homogeneity of variance tests performed on the scaled MSE matrices reveal a pattern not seen in the MSE matrices (S1 Appendix). With scaled MSE, all methods utilizing feature-fixing have lower variance than their counterparts without feature-fixing. The variance of Viterbi methods utilizing feature-fixing is similar to the variance of non-Viterbi methods without feature-fixing. In general, scaled MSE distances are substantially reduced since feature-fixing tends to increase pairwise cP distances but not landmark MSEs. This effect can also be seen in the matrix heat maps (S2 Appendix), and supports the significance of feature-fixing as a important parameter for the performance of the tree-based improvement methods.

### Quantitative comparison of ordinations generated by globally informed methods

The previous analyses permit the identification of those methodological parameters with strong effects on shape space characterization, but they are not informative regarding certain aspects (e.g., is feature-fixing beneficial?). Our final analysis compares the ordinated shape spaces of those globally informed methods that are most and least similar to the shape space characterized by user-determined landmarks. Principal components analysis of the vectors encoding method parameter combinations (S1 Appendix) reveals these methods (Fig 4), which are subsequently referred to as GLobal-informed Automated Method 1 (GLAM1: LAST; α = balance; no feature-fixing), GLAM2 (LAST; composed; α = balance; no feature-fixing), and GLAM3 (LAST; composed; α = balance; feature-fixing). In addition to these three novel methods, we also compare the shape spaces generated by user-determined landmarks and auto3Dgm.

**Fig 4.**
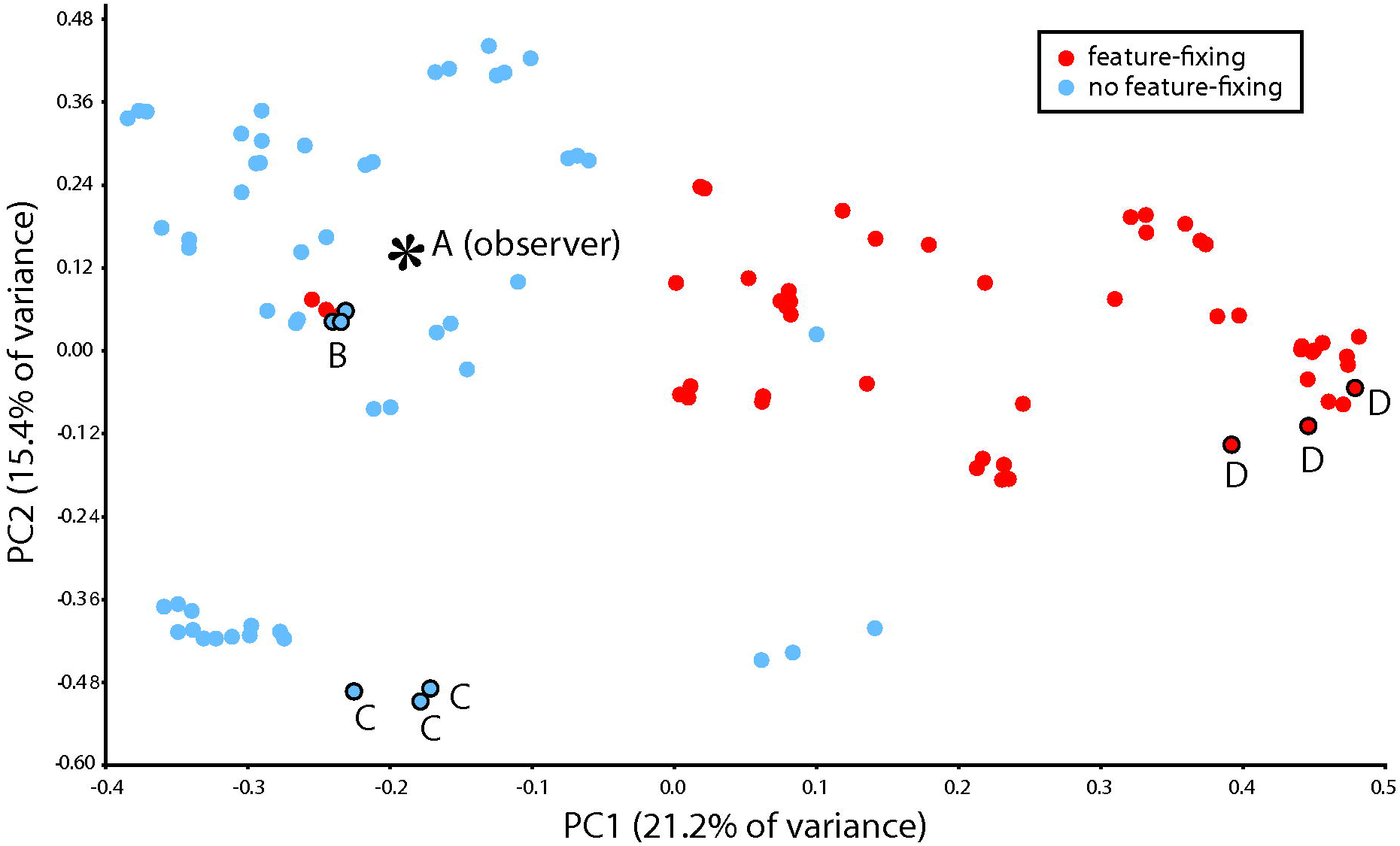
Results of principal component analysis comparing different parameter combinations of automated alignment and mapping. Each point represents an automated analysis of 116 tooth surfaces from [1]. Points that plot close together represent analytic protocols yielding similar representations of shape affinities for the surfaces in the test dataset. Note that only around 36% of the total variance is represented on these first two principal components. As confirmed by the statistical analyses detailed in the text, this plot indicates feature-fixing has the strongest effect on shape affinities among sampled teeth. To our surprise, analyses without feature-fixing characterize shape affinities in a way more similar to the user-based “ground truth”. Three treatments are examined in more detail: A) user-determined landmarks; B) GLAM1 (LAST; = balance; no feature-fixing), the approach most similar to the “ground-truth”; C) GLAM2 (LAST; composed; α = balance; no feature-fixing); and D) GLAM3 (LAST; composed; α = balance; feature-fixing). The separation between A and both C and D suggest the latter two approaches characterize shape affinities in a distinct manner relative to A. As each of these treatments was run with three different pseudolandmark resolutions, each treatment is represented three times. The relatively minor variance of each treatment under differing resolutions demonstrates that pseudolandmark sampling has little effect on shape space characterization (also confirmed by statistical analyses in the text).

In order to compare the ordinated shape spaces of these five methods, we first identified 18 phylogenetically cohesive taxonomic groups (S1 Appendix) that were fairly distinctive when visualized on the first two principal components of the user-determined landmarks (Fig 5a). We then ran one-way ANOVAs on a vector representing the first 46 principal component scores (S1 Appendix). As might be expected from the taxonomic separation apparent in Fig 5, we found all samples to have highly significant interspecific variance (Table 9), with the highest significance level for the user-determined landmarks. Treatments more similar to the user-determined landmarks in Fig 5 (GLAM1 and GLAM2) had higher significance levels than the treatments farther removed from the ground truth (GLAM3). Though auto3Dgm does not specify explicit maps relating surfaces, its ordinated shape space appears to be most similar to the user-based result in this limited (though potentially representative) analysis.

**Fig 5.**
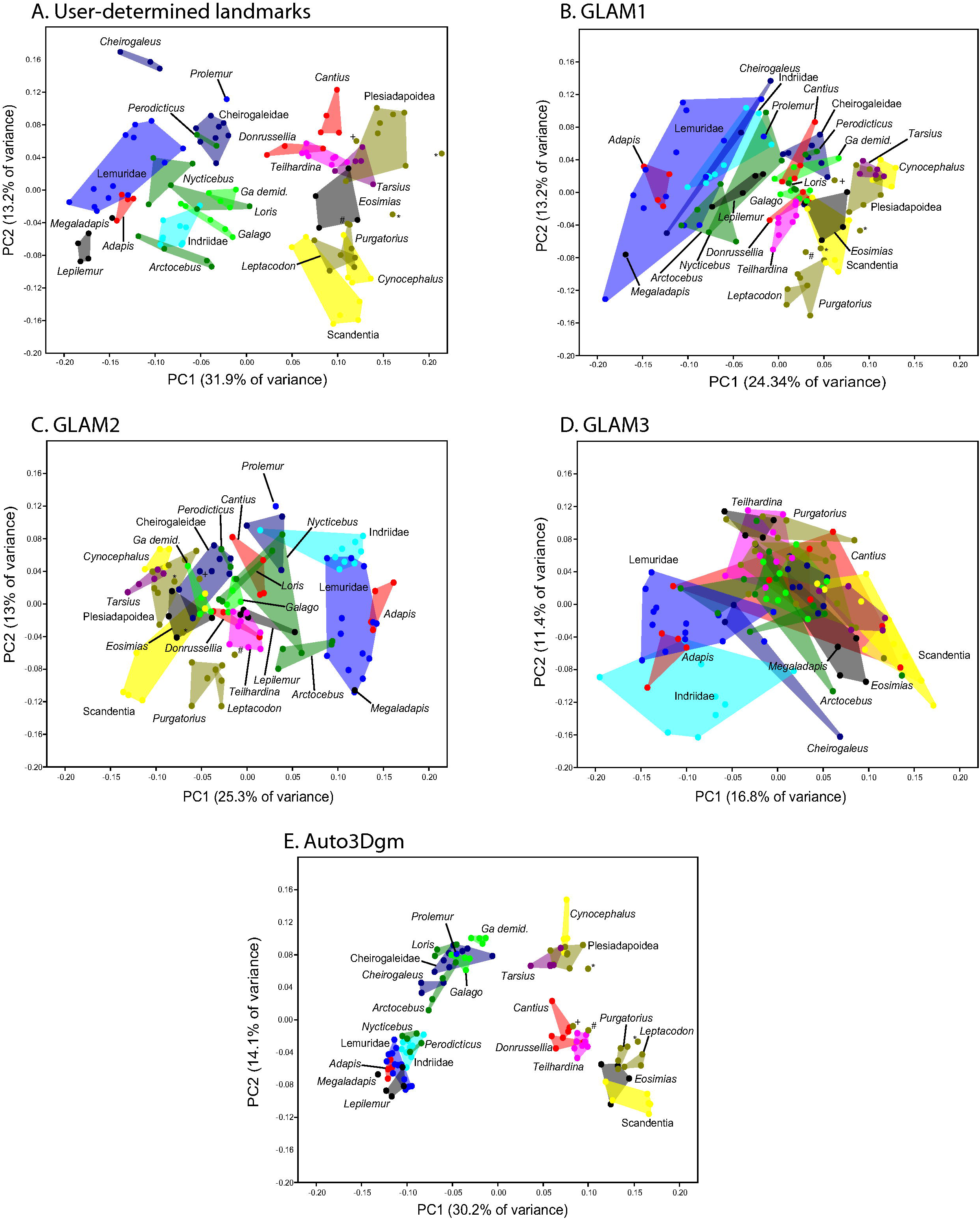
Principal components analyses characterizing shape affinities in a sample of 116 teeth using five alternative approaches. Approaches include: A) User-determined landmarks; B) GLAM1 (LAST; α = balance; no feature-fixing), the approach most similar to the “ground-truth” in Fig 4; C) GLAM2 (LAST; composed; α = balance; no feature-fixing); D) GLAM3 (LAST; composed; α = balance; feature-fixing); and E) auto3Dgm. Neither C nor D was expected to look similar to A or B based on Fig 4. Minimum convex polygons include individual specimens of closely related species expected to be similar based on visual inspection and traditional comparative analyses. The degree to which each method successfully distinguished groups was evaluated with a series of vector ANOVAs in which taxonomic groups shown in these images were the treatment effects. From these analyses it appears that A, B, and E do the best job of separating taxonomic groups (Table 9). The specific surfaces used in each group and the data for each analysis is provided (S1 Appendix). Exemplar teeth of each group are shown in Fig 7.

**Table 9.**
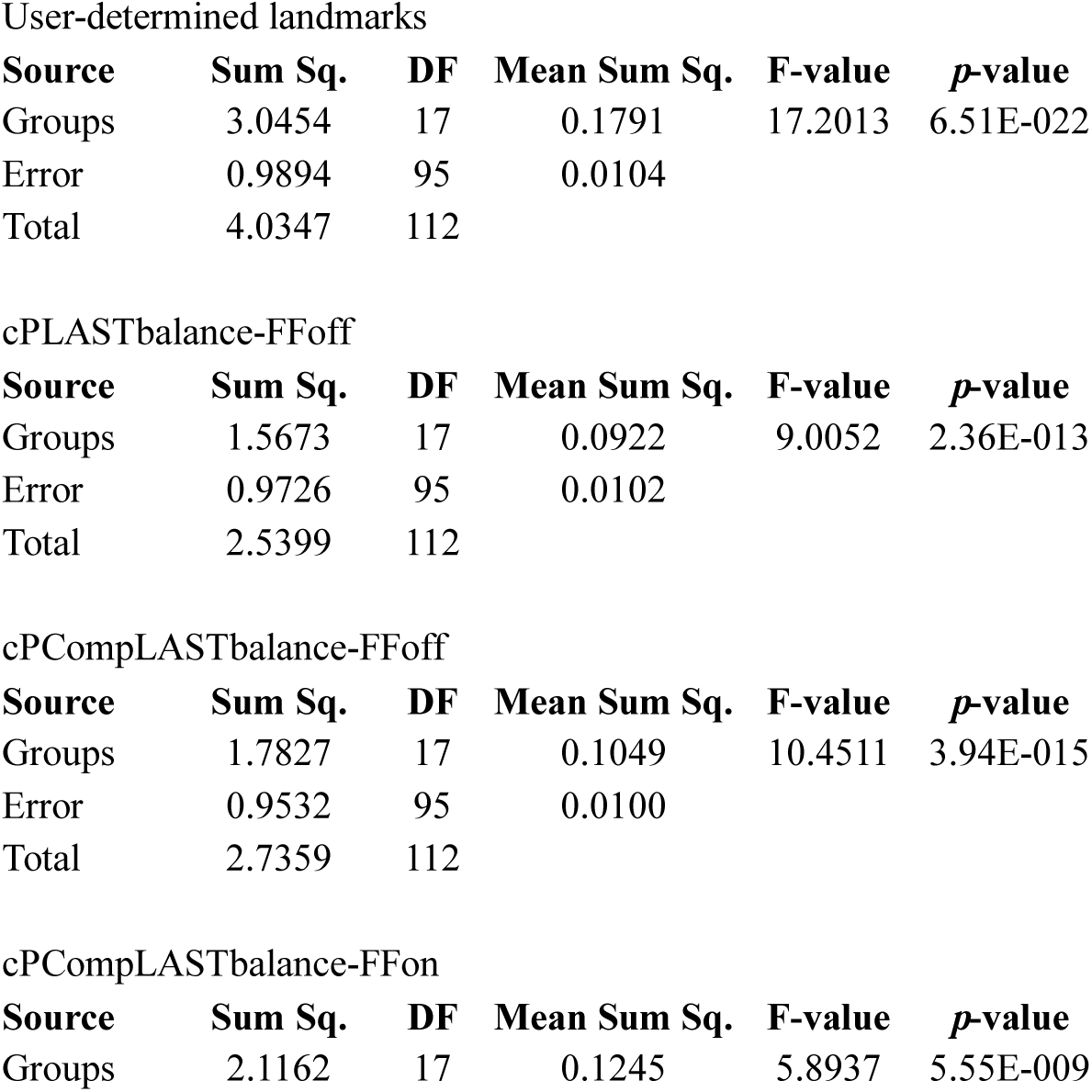
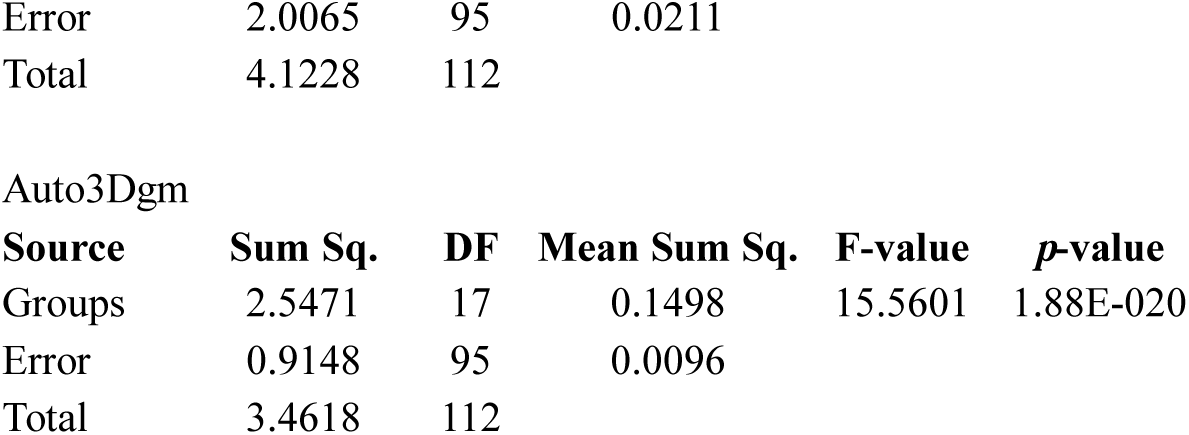
One-way ANOVA on taxomonic groups to assess which parameter combinations result in shape spaces with greatest between-group distinctiveness.

User-determined landmarks
| Source | Sum Sq. | DF | Mean Sum Sq. | F-value | p-value |
| --- | --- | --- | --- | --- | --- |
| Groups | 3.0454 | 17 | 0.1791 | 17.2013 | 6.51E-022 |
| Error | 0.9894 | 95 | 0.0104 |  |  |
| Total | 4.0347 | 112 |  |  |  |

cPLASTbalance-FFoff
| Source | Sum Sq. | DF | Mean Sum Sq. | F-value | p-value |
| --- | --- | --- | --- | --- | --- |
| Groups | 1.5673 | 17 | 0.0922 | 9.0052 | 2.36E-013 |
| Error | 0.9726 | 95 | 0.0102 |  |  |
| Total | 2.5399 | 112 |  |  |  |

cPCompLASTbalance-FFoff
| Source | Sum Sq. | DF | Mean Sum Sq. | F-value | p-value |
| --- | --- | --- | --- | --- | --- |
| Groups | 1.7827 | 17 | 0.1049 | 10.4511 | 3.94E-015 |
| Error | 0.9532 | 95 | 0.0100 |  |  |
| Total | 2.7359 | 112 |  |  |  |

cPCompLASTbalance-FFon
| Source | Sum Sq. | DF | Mean Sum Sq. | F-value | p-value |
| --- | --- | --- | --- | --- | --- |
| Groups | 2.1162 | 17 | 0.1245 | 5.8937 | 5.55E-009 |

|  |  |  |  |
| --- | --- | --- | --- |
| Error | 2.0065 | 95 | 0.0211 |
| Total | 4.1228 | 112 |  |

Auto3Dgm
| Source | Sum Sq. | DF | Mean Sum Sq. | F-value | p-value |
| --- | --- | --- | --- | --- | --- |
| Groups | 2.5471 | 17 | 0.1498 | 15.5601 | 1.88E-020 |
| Error | 0.9148 | 95 | 0.0096 |  |  |
| Total | 3.4618 | 112 |  |  |  |

User-determined landmarks recover substantially more between-taxon variance than globally informed methods (Table 9). The increase in between-group variance suggests that user-determined landmarks are better at capturing “real” between-taxon differences. It is quite likely that user-determined landmarks are focused on those features that exhibit a large amount of between-group variance, so that morphological expertise permits more variance to be captured by relatively fewer landmarks. This focus has been argued to be the primary advantage that semiautomated methods have over fully automated alternatives [52]. However, since the manual landmarks were collected with a *priori* taxonomic knowledge, it is also possible the researcher inadvertently biased their landmarking to maintain intraspecific consistency. Repeating manual landmark data collection would mitigate such bias and insure the absence of such effects. However, to avoid adding more sources of error, any additional data collection would have to be done by a researcher with equivalent anatomical expertise, which risks introducing a similar bias if they remember species-specific morphological patterns well. A possible approach would be to collect landmark data in random taxonomic order over a widely spaced time interval.

Finally, we were surprised to find that feature-fixing generally reduced the similarity between globally informed methods and user-determined landmarks (Fig 4). Since user-determined landmarks are often close to (but not necessarily exactly coincident with) positions of extreme geometric configuration, the discrepancy between user-determined and feature-fixed landmarks can be as large as several edges away on the discretized triangular mesh. Such a difference is comparable to the magnitude of some landmark MSEs between very similar shapes. Additionally, the TPS procedure does not control for shape distortion at regions lacking anchor points, though some user-determined landmarks are positioned in such regions. Therefore, an alternative approach to TPS, with guaranteed low distortion on the regions of shapes even without geometric characteristics, may be preferable for geometric morphological analysis.

### Qualitative assessment and biological implications of ordinations generated by globally informed methods

In order to understand how geometric affinities of particular taxonomic groups differ qualitatively across methods, we now focus on the details exhibited by each ordination. Fig 6 provides terminology for some notable features of therian (marsupial and placental mammals) mandibular molars, and Fig 7 provides examples of these teeth for the sample used in this study. For all methods, the presence and prominence of the paraconid, the most anterior cusp of the tooth, drives variation on PC1 (Fig 5). User-determined landmarks and auto3Dgm are most similar in this pattern: for these two methods, there is a slight gap between the distributions of extant strepsirrhines (all lack a paraconid) plus *Adapis* (which has a relatively small paraconid) and tarsiers, non-primates, and the remaining fossil taxa (all of which have prominent paraconids) (Fig 7). This distinction blurs in the ordinations of globally informed methods as those strepsirrhines with relatively prominent trigonids invade the space of paraconid-bearing taxa. Specifically, lorisiforms (galagos, *Nycticebus, Perodicticus, Loris* and *Arctocebus), Lepilemur*, and all cheirogaleids except *Cheirogaleus* (which has a strongly reduced trigonid) overlap with early fossil euprimates *(Teilhardina, Cantius*, and *Donrussellia)*, certain plesiadapiforms (e.g. *Pronothodectes)*, and the treeshrew *Ptilocercus* in the three globally informed methods. Among these methods, the overlap is least pronounced in GLAM1, which also happens to be the globally informed method that is closest to user-determined landmarks in Fig 4.

**Fig 6.**
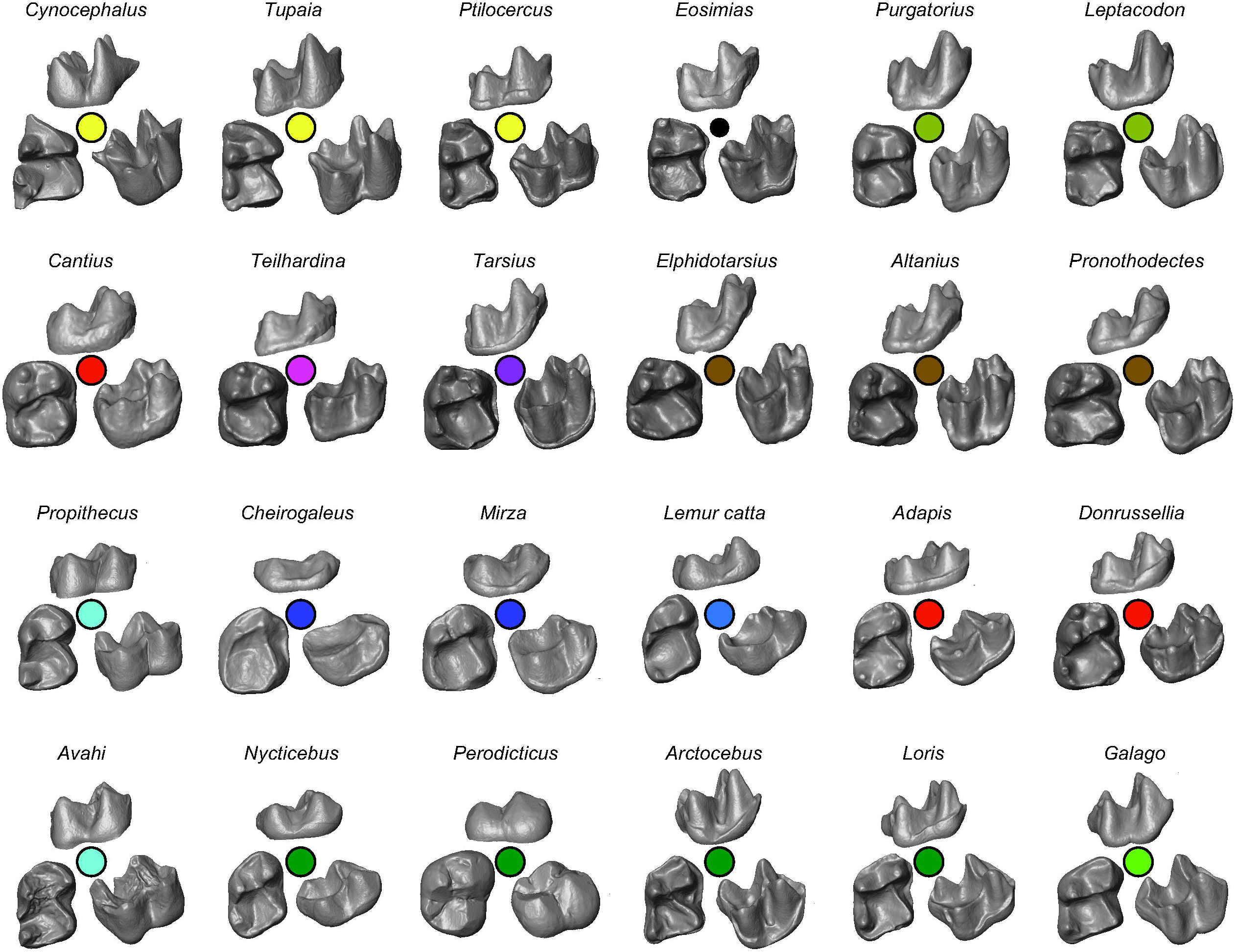
Nomenclature of the primary features of a therian mandibular molar. Occlusal view.

**Fig 7.**
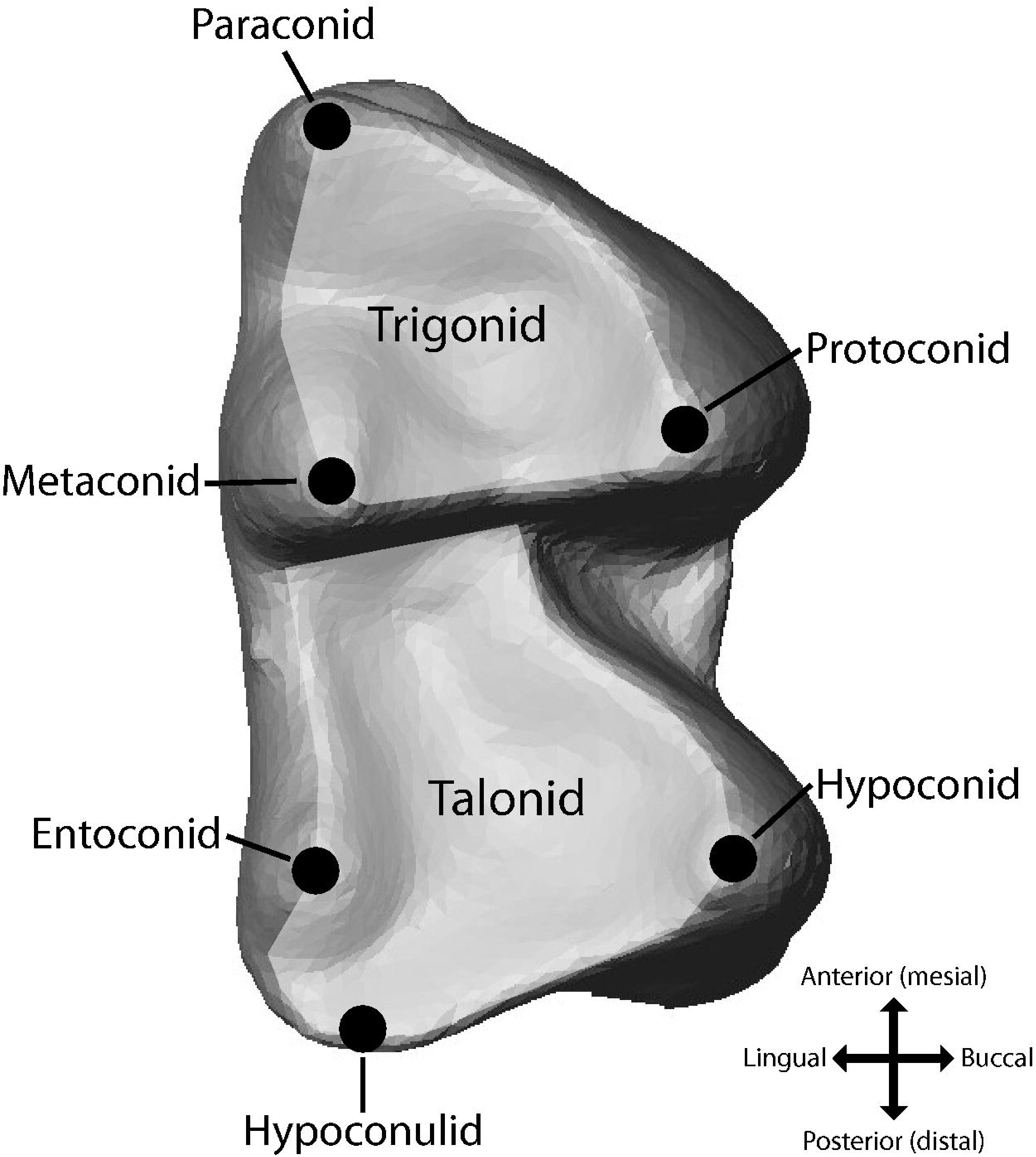
Images of teeth of representative taxa in this study. For each taxon, views are buccal (upper), occlusal (lower left), and a three-quarter profile (lower right). Colors correspond to minimum convex polygons shown in Fig 5.

PC2 of the user-determined landmark plot (Fig 5a) separates teeth with a hypoconulid close to the entoconid and substantially posterior to the hypoconid (e.g., *Lepilemur* and *Tupaia)* from teeth with a more mesially and buccally positioned hypoconulid (e.g., *Cheirogaleus).* Trends of variation in PC2 of some globally informed methods are similar to the user-determined landmarks in certain respects. However, none of the automated 3DGM methods reflect variation in the talonid cusps on PC2, as indicated by the different positions of *Cynocephalus*, galagos, and certain lorisids (these taxa all have strongly lingually positioned hypoconulid). The auto3Dgm plot, with the smallest within-group distributions, is most similar to the hypoconulid-driven trend of the user-determined landmarks, but differs in the recovered overlap of galagos and cheirogaleids. In contrast, the user-determined landmarks pull galagos toward lemurids and indriids. Likewise, the interspecific distribution of lorisids is inverted between the user-based approach and auto3Dgm. With user-determined landmarks, *Arctocebus* and *Loris* overlap more with indriids and lemurids, while *Perodicticus* and *Nycticebus* overlap with cheirogaleids. In auto3Dgm, the trend is reversed (though galagos overlap most extensively with *Arctocebus* and *Loris* in both methods). In fact, auto3Dgm shows tight clustering of these taxa, revealing strong affinities of *Nycticebus* and *Perodicticus* with indriids and lemurids, and affinities of *Loris* and *Arctocebus* with cheirogaleids and galagos. From a phylogenetic perspective, these alternative groupings are not intuitive. However, from a functional perspective, the user-determined landmarks group frugivorous lorisids (e.g., *Perodicticus)* and omnivorous cheirogaleids (e.g., *Microcebus)* in one region and insectivorous lorisiforms (e.g., *Arctocebus* and galagos) with folivorous indriids (e.g., *Avahi)* in another.

The globally informed method most similar to the user-based approach in Fig 4 (GLAM1, Fig 5b) does not exhibit any obvious trends on PC2 for strepsirrhines, but shows a distribution of the four lorisids more consistent with a dietary interpretation (low relief frugivorous lorises overlapping with omnivorous cheirogaleids, and high relief insectivorous lorisiforms overlap with insectivorous galagids and folivorous indriids). In other automated methods, taxonomic groups overlap too extensively for succinct description, including among lorisids. Nonetheless, all three of the globally informed methods (Fig 5b-d) preserve separation between galagos and cheirogaleids (like user-determined landmarks but unlike auto3Dgm) while also maintaining a large separation between galagos and indriids (unlike user-determined landmarks). Therefore, from a phylogenetic perspective, the globally informed methods return more intuitive results than either user-determined landmarks or auto3Dgm.

The relative positions of non-primate taxa and primitive fossil primates are similar in the five example plots (Fig 5). The extreme values are typically tupaiid treeshrews (an extant non-primate that is insectivorous), and tend to cluster close to *Leptacodon* (a fossil non-primate) and *Purgatorius* (the oldest and most basal known stem-primate). *Ptilocercus*, a more omnivorous treeshrew, is typically separated from this cluster. The ambiguous fossil taxon *Altanius* (either euprimate or stem-primate) also plots near this cluster but typically has less extreme PC2 values. Eosimiidae, a group of purported stem-anthropoid primates, also plots near the tupaiid-*Leptacodon-Purgatorius* cluster with at least one individual always plotting close to *Ptilocercus.* Early euprimates *Cantius, Teilhardina*, and *Donrussellia* are near one another with slightly less extreme PC1 and PC2 scores. Stem-primates more derived than *Purgatorius (Paromomys, Plesiolestes, Pronothodectes, Chronolestes, Elphidotarsius)* tend to plot separately from the early euprimates (but further from treeshrews, *Purgatorius*, Eosimiidae, *Altanius*, etc.) while overlapping *Tarsius* (an extant haplorrhine).

Finally, the non-primate *Cynocephalus* plots with treeshrews in the user-based method, a result consistent with these taxa being outgroups to primates. Similarities between *Cynocephalus* and *Tupaia* in the user-based result are driven by the strongly lingual and posterior position of the hypoconulid. In contrast, all of the globally informed methods show *Cynocephalus* overlapping primarily with *Tarsius*, linking a dermopteran to a crown haplorhine. The automated results seem to reflect gross similarities between *Cynocephalus* and *Tarsius* (e.g., both taxa have relatively square occlusal outlines), but they are not phylogenetically intuitive.

Of the methods compared, GLAM3, the only method utilizing feature-fixing, shows the least taxonomic differentiation (Fig 5d). It is also the only method that does not place the subfossil *Megaladapis* in the region occupied by *Adapis* and lemurids. The early euprimate *Cantius* is scattered across the entire plot area. Stem-primates, *Leptacodon, Purgatorius*, and *Teilhardina* are largely overlapping and oddly plot near *Lepilemur* (Fig 5d). *Cheirogaleus* spans over half the range of PC2 values. This scatter is reflected in the relatively low value obtained by taxonomic ANOVA for this group as well (Table 9).

Two observations suggest that GLAM1 and GLAM2 (which do not use feature-fixing) reflect sample geometry better than GLAM3 (which uses feature-fixing). First, in the plot of method by treatment type (Fig 4), they were closer to the result from user-determined landmarks. Second, the one-way ANOVA on taxonomic groups of the vector distribution of ordinations produced by GLAM1 has a higher p-value than GLAM3, suggesting greater taxonomic distinctiveness (Table 9). It was surprising that auto3Dgm has a pattern and magnitude of taxonomic distinctiveness more similar to the user-determined landmarks; in many ways, auto3Dgm is relatively naïve compared to other tree-based techniques presented in this paper.

The observation that feature-fixing degrades taxonomic signal is worth further investigation. TPS, the main technique involved in feature-fixing, strives to align specified corresponding anchor points and generate a smooth interpolation between the shapes without guaranteed control for the distortion of the final map. The cP distance, in a certain sense, measures the minimum global average distortion of a class of candidate maps between two surfaces. Therefore, maintaining low global distortion may be more important for producing cP distances that faithfully reflect the geometric dissimilarity than precisely matching geometrically characteristic point features. Furthermore, the quality of the interpolated map depends heavily on the choice of anchor points: if two corresponding anchors appear to close to each other, TPS will face numerical stability issues. If the correspondences between anchor points are incomplete or wrongly specified (e.g. for teeth of low relief and probably fewer detectable extremal points such as the omnivorous *Cheirogaleus)*, TPS may generate less biologically meaningful maps. It is therefore possible that the various sources of map and distance distortion in feature-fixing methods generate less taxonomically cohesive results. Rather than a major improvement for the automated 3DGM methods, feature-fixing may primarily facilitate map visualization compared to existing geometric morphological analysis.

## Conclusions

In this study, we have addressed and overcome limitations of previously published automated 3DGM methods, cPDist and auto3Dgm, [1,2] and provided a detailed description of how the results of existing and novel automated methods compare to a user-determined landmark approach. Both the globally informed methods proposed here and auto3Dgm reflect the similar geometric patterns as user-based methods. Relative to cPDist, the dramatic reduction in MSE of propagated landmarks of the globally informed methods shows that global sample information is a critical component of automated analysis on samples with large shape differences. Other modifications did not definitively improve the similarity to a user-based approach. In particular, feature-fixing, or the automatic manipulation of maps to maintain type II landmark [53] representation, appears to add error and reduce taxonomic cohesiveness. Though composed LAST methods may potentially reduce the increased map inaccuracy between similar shapes that are not directly connected by an edge in the MST, no approach permits full retention of map quality of direct comparisons between similar shapes. This suggests that developing alternative approaches for analyzing collections of highly dissimilar shapes remains an interesting and challenging problem.

### Comparison of novel globally informed methods and auto3Dgm

We were surprised that auto3Dgm produced ordinations with cohesive taxonomic groups that were also phylogenetically and geometrically intuitive given our understanding of feature variation in the sample (Fig 5e, Table 9). Though it is not entirely clear that tight taxonomic clustering is the most accurate expression of shape differences in the sample, the recovered pattern does lead to the question of whether or not any of the novel methods presented here are superior to auto3Dgm in regard to their characterization of shape affinities. We believe the novel globally informed methods represent improvements for three reasons: 1) pseudolandmark sampling density does not appear to affect the novel methods of this study, 2) the novel methods produce more intuitive ordinations, and 3) the novel methods evenly fill the ordination space. First, Vitek et al. [29] found that ordinations produced by auto3Dgm are sensitive to pseudolandmark sampling density (i.e., number of points per tooth). In this study, because downstream analyses were not sensitive to sampling density, this does not appear to be a problem for the methods presented in this study. Second, several inter-taxonomic affinities in the auto3Dgm result differ strongly from user-determined landmarks and are less functionally or geometrically intuitive than results from the novel methods of this study. In the case of the relative positions of galagos, cheirogaleids, and indriids, results from the novel methods are more intuitive, in both a functional and phylogenetic sense.

Finally, the novel methods evenly fill the ordination space in a manner more similar to the user-based approach (Fig 5). The distribution of specimens in the auto3Dgm ordinations often form a Y pattern in which points cluster linearly through regions of space, giving a much different perspective on how filled the shape space is compared to user-based methods (Fig 5). The Y pattern also seems to appear in interspecific PCA ordinations generated by Generalized Procrustes Surface Analysis [54], a recent shape analysis method of similarly high dimensionality. The diffuse distribution seems problematic, as it may reflect a highly skewed distribution of values in the correlation matrix of the PCA. Gonzalez et al. [27] recommend converting pseudolandmarks to sliding landmarks to minimize surface bending energy or average Procrustes distance between specimens. In this way, they eliminate the Y pattern; however, this may potentially reflect the addition of random noise. Alternatively, replacing PCA with a dimension reduction method more suitable for high dimensional data (such as t-distributed stochastic neighbor embedding [55]), may reduce the strength of the Y pattern. Whatever the remedy for auto3Dgm, the Y pattern does not manifest in any of the novel methods presented here, suggesting they are indeed improvements to existing automated methods.

### Optimal applications for automated approaches

Because the examined parameter combinations of globally informed methods and auto3Dgm produce similar shape ordinations that are largely consistent with results of a user-based approach, we recommend using automated approaches when: 1) the primary questions concern patterns of overall variation in biological shapes, 2) a large number of type II landmarks (defined in [53] as areas of high local curvature) are not consistently available, or 3) measurement/landmark selection may increase the potential for biased results. These recommendations are similar to those of Gonzalez et al. [27], who evaluated the performance of auto3Dgm against a semi-automated landmark method. The first two conditions are related, since fewer type II landmarks limit the ability to assess overall variation. User-based approaches for assessing shape *disparity* (which refers to the overall structure of interest) become increasingly limited as sample diversity increases. Polly [56] highlights the limitation of user-based approaches that require many biologically equivalent landmarks to represent patterns of shape variation and emphasizes the importance of methods that lack this requirement. How can one represent the absence of a paraconid quantitatively if the feature does not exist in all specimens? The researcher could landmark the space where the paraconid “would be”, but this is obviously subjective. Automated 3DGM methods do not make such explicit assumptions about feature equivalence and therefore can more comprehensively measure shape variation.

The third scenario, reducing the potential for user bias, is particularly salient when the anatomical structure of a fossil has been linked to a particular taxonomic identification or phylogenetic hypothesis. In these cases, it is critical that the researcher’s qualitative assessments match quantitative approaches and that selective choice of measurements and/or landmarks has not unfairly weighted the evidence toward a particular hypothesis. Boyer et al. [24] used an automated analysis for this role: a newly discovered fossil was shown objectively to be a range extension of a previously unknown species. In the dataset considered in this paper, the family Eosimiidae has controversial affinities. The group is typically assigned to anthropoid primates [57–61]), but this assessment has been questioned [62–66]. Even if the diagnosis as an anthropoid relative is accepted, it is unclear what the geometry of eosimiid tooth structure might reveal about initial stages of anthropoid evolution.

Given these issues, our more quantitative, more comprehensive, and more objective assessment of eosimiid tooth form should interest anthropologists and paleontologists, as results show the second mandibular molar of eosimiids is distinctively more similar to non-primates and stem-primates of the sample than to early euprimates or tarsiers. This result is recovered by all methods compared here, including user-determined landmarks. It is consistent with the perspective that Eosimiidae may be positioned more basally than stem Anthropoidea *(contra* [57–59]), and questions purported similarities between Eosimiidae and Tarsiidae. Furthermore, the recovered dental affinities are consistent with recent results from analyses of Eosimiidae ankle bones [31,67,68] that suggest eosimiid ankle morphology is similar to that inferred for the common ancestor of primates of modern aspect. In our opinion, these findings point to the need for continued examination of eosimiid relationships, anthropoid relationships, and certain patterns of early primate evolution.

In sum, this study affirms the utility and reliability of automated approaches through thorough comparisons of automated approaches and user-based approaches. The globally informed methods presented here have several important advantages over user-based approaches and previously published automated 3DGM methods. Future work will improve the use of global sample information in computing correspondence maps between objects and will hopefully recover even greater geometric fidelity in automated morphometric analysis.

## Supporting Information

**S1 Appendix. Supplementary tables.** Includes principal components scores for the method ANOVA, results of MSE variance tests, MRPP results, groups included in taxonomic ANOVAs, and principal component scores for taxonomic ANOVAs (XLS).

**S2 Appendix. Supplementary figures and code.** Includes matrix heat maps, bivariate plots of landmark MSES for globally informed methods, MSE variance boxplots, explanations of ANOVA and linear mixed model equivalents, and MATLAB code for the vector equivalents (PDF).

## Acknowledgments

Our thanks to curators and staff at the American Museum of Natural History for providing access to dental specimens to be molded, cast, and μCT-scanned; to D. Krause and J. Groenke of Stony Brook University for access to facilities at the Vertebrate Paleontology Fossil Preparation Lab; to C. Rubin and S. Judex of Stony Brook University for access to μCT scanners; and to K. Christopher Beard for access to eosimiid dental casts. Grant support provided by NSF BCS 1304045 (to DMB and EM St. Clair), NSF BCS 1317525 (to DMB and ER Seiffert), NSF BCS 1440742 (to DMB and GF Gunnell), NSF BCS 1440588 (to JI Bloch), and NYCEP IGERT (NSF DGE-0991660).

## Footnotes

1. Note that when local distance distortion is measured by the ratio between energy of the composed map and the direct distance (i.e. energy of the direct map), it is possible for a =1 even when the two maps are different. This is in contrast with measuring local distance distortion by the ratio between the length of the minimum path and the direct distance, in which case a =1 requires the minimum path to equal the direct link and thus forcing the entire tree to be a SPT.

